# Single-cell and spatial transcriptomics of vulvar lichen sclerosus reveal multi-compartmental alterations in gene expression and signaling cross-talk

**DOI:** 10.1101/2024.08.14.607986

**Authors:** Peng Sun, Christina N. Kraus, Wei Zhao, Jiahui Xu, Susie Suh, Quy Nguyen, Yunlong Jia, Arjun Nair, Melanie Oakes, Roberto Tinoco, Jessica Shiu, Bryan Sun, Ashley Elsensohn, Scott X. Atwood, Qing Nie, Xing Dai

**Author notes:** Correspondence: C.K and X.D. These authors contributed equally (Peng Sun, Christina N. Kraus).

## Abstract

Vulvar diseases are a critical yet often neglected area of women’s health, profoundly affecting patients’ quality of life and frequently resulting in long-term physical and psychological challenges. Lichen sclerosus (LS) is a chronic inflammatory skin disorder that predominantly affects the vulva, leading to severe itching, pain, scarring, and an increased risk of malignancy. Despite its profound impact on affected individuals, the molecular pathogenesis of vulvar LS (VLS) is not well understood, hindering the development of FDA-approved therapies. Here, we utilize single-cell and spatial transcriptomics to analyze lesional and non-lesional skin from VLS patients, as well as healthy control vulvar skin. Our findings demonstrate histologic, cellular, and molecular heterogeneities within VLS, yet highlight unifying molecular changes across keratinocytes, fibroblasts, immune cells, and melanocytes in lesional skin. They reveal cellular stress and damage in fibroblasts and keratinocytes, enhanced T cell activation and cytotoxicity, aberrant cell-cell signaling, and increased activation of the IFN, JAK/STAT, and p53 pathways in specific cell types. Using both monolayer and organotypic culture models, we also demonstrate that knockdown of select genes, which are downregulated in VLS lesional keratinocytes, partially recapitulates VLS-like stress-associated changes. Collectively, these data provide novel insights into the pathogenesis of VLS, identifying potential biomarkers and therapeutic targets for future research.

## INTRODUCTION

Lichen sclerosus (LS) is a chronic inflammatory skin disease that predominantly affects the vulva, leading to pruritus, pain, irreversible scarring, decreased quality of life, and an increased risk of malignant transformation to squamous cell carcinoma^1,2^. LS also affects extragenital skin and penile skin, though less commonly^3^. Vulvar LS (VLS) has been estimated to affect up to 1.7% of patients presenting to general gynecology clinics^4^. However, LS is likely underdiagnosed, making true incidence and prevalence challenging to estimate^5^. Histopathologically, VLS is often characterized by a lichenoid tissue reaction combined with dermal collagen changes, with varying degrees of epidermal alterations and dermal sclerosis or fibrosis^6,7^. Despite descriptions of the stages of LS in dermatopathology textbooks, there is a notable lack of comprehensive studies. Current literature primarily references individual studies that present similar findings, such as the progression of lymphocytic infiltrate from the dermo-epidermal junction (DEJ) through the dermis^8^. The histopathological variations, coupled with overlapping features with other vulvar conditions and treatment-induced changes complicate the accurate diagnosis and histological interpretation of VLS^6,9^. Currently, there are no FDA-approved therapies for VLS. The standard of care involves the use of ultrapotent topical steroids, with evidence suggesting lifelong maintenance use is required to prevent scarring and malignancy^1,10,11^. However, some LS cases are recalcitrant to topical steroids and overuse can exacerbate the atrophy associated with this condition. Additionally, many patients suffer from steroid phobia^12^, which hinders the effective use of medication to treat the disease and prevent long-term complications. While circumcision is usually curative for LS in penile skin^13^, managing VLS is challenging due to the lack of evidence-based therapeutic alternatives, as well as underreporting and stigmatization that often lead to advanced disease stages.

Despite its significant morbidity, the pathogenesis of LS is poorly understood, hindering the development of effective therapies. It is believed that both environmental factors (such as trauma) and genetic predisposition play a role in the development of LS, with a positive family history reported in 12% of individuals and an increased association with HLA alleles, including HLA-DQ7^9,14^. Some consider LS to be an autoimmune condition, but this classification remains controversial^9,15^. Others argue that LS, particularly in males, is caused by external stressors such as urine^16^.Dense infiltration of lymphocytes composed of CD4^+^, CD8^+^, and regulatory T cells can be found at the DEJ or in the dermis of LS skin^9,15^. Expression of chemokine receptors CXCR3 and CCR5, coupled with a lack of CCR3 and CCR4 expression, is thought to suggest a T helper (Th1) response mediated by IFN-γ production, though Th2 cytokine production has also been reported^17,18^. Other reported alterations include presence of autoantibodies against ECM1 in the serum of LS patients^18^, and increased synthesis of collagen I, collagen III, galectin-7, and microRNA-155 in LS samples^3,19^. Involvement of oxidative stress was also suggested^20^. Despite these tantalizing clues, existing studies on LS pathogenesis are limited in depth and scope, focusing on a small number of cellular or molecular markers. To date, the most comprehensive study is bulk RNA sequencing analysis of VLS lesions suggestive of a T cell response^21^. As such, the specific molecular alterations in different cell populations of the VLS skin have not been characterized. Furthermore, a comprehensive understanding of cell-cell communications that orchestrates VLS inflammation and pathology is lacking.

Single-cell RNA-sequencing (scRNA-seq) has greatly expanded our ability to gain novel insights into the mechanisms of various inflammatory skin diseases by dissecting individual cell types, states, and intercellular interactions. Concurrently, spatial transcriptomics offers insights into the localization of cell types and states, and the dynamics of cell-cell communications at disease sites. Here, we employ these techniques to achieve a comprehensive view of the cellular and molecular differences between non-lesional (NL) and lesional (LE) skin of VLS patients. Our findings reveal gene expression alterations in fibroblasts, immune cells, keratinocytes, and melanocytes, along with a systematic overview of signaling communication changes in LE skin. We also provide evidence for keratinocyte apoptosis and necroptosis as possible mechanisms of cell death in LE skin. Finally, we demonstrate that the siRNA-mediated knockdown of specific genes, identified by our scRNA-seq analysis as downregulated, in human keratinocytes partially recapitulates the gene expression changes observed in LE keratinocytes. Specifically, knockdown of *ATF3* or *APOE* induces a keratinocyte stress response observed in LE epidermis. In summary, this study offers the first comprehensive characterization of VLS skin lesions, revealing new insights into the multi-compartmental nature of VLS pathogenesis and serving as a vital resource for vulvar skin disease research.

## RESULTS

### LE skin from VLS patients exhibits inter-patient heterogeneity in immune infiltration, dermal and epidermal alterations, and cell death

To enhance morphological and biochemical understanding of VLS skin, we conducted histologic analysis coupled with immunostaining on clinically archived formalin-fixed paraffin-embedded (FFPE) human vulvar tissue specimens. We compared LE samples from VLS patients at different clinical stages of the disease to NL samples from the same patients and to samples from healthy controls (HC) without VLS. As reported by others^22^, we observed remarkable intra- and inter-patient heterogeneity in the histological features of VLS lesions. These ranged from predominantly inflammatory or sclerotic subtypes, each accompanied by varying degrees of epidermal changes (Fig. 1A; Table S1). Sclerotic changes, characterized by marked dermal collagen homogenization, appeared to start in the papillary dermis and subsequently spread to the reticular dermis, leaving behind a largely acellular extracellular matrix (Fig. 1A4, A4’). Immunostaining using fibroblast intermediate filament protein vimentin and pericyte/smooth muscle cell marker SMA did not reveal a clear alteration in the dermis of non-sclerotic LE relative to NL (Fig. S1A). Consistent with the varying degrees of lichenoid inflammation, immunofluorescence using CD45 (a marker for all immune cells) and CD3 (a marker for T cells) revealed differing levels of total immune cell and T cell abundance in both the epidermis and dermis (Fig. 1B; S1B). Additionally, immune cells were present in cellular aggregates deep in the reticular dermis of fully sclerotic LE skin (Fig. S1B). Interestingly, while immune cells were also detected in NL and HC skin, they appeared to adopt a more elongated or dendritic morphology, while those in LE skin had a more rounded shape (Fig. 1B; S1B, C), possibly reflecting enrichment of different types of immune cells in HS/NL versus LE skin.

**Figure 1.**
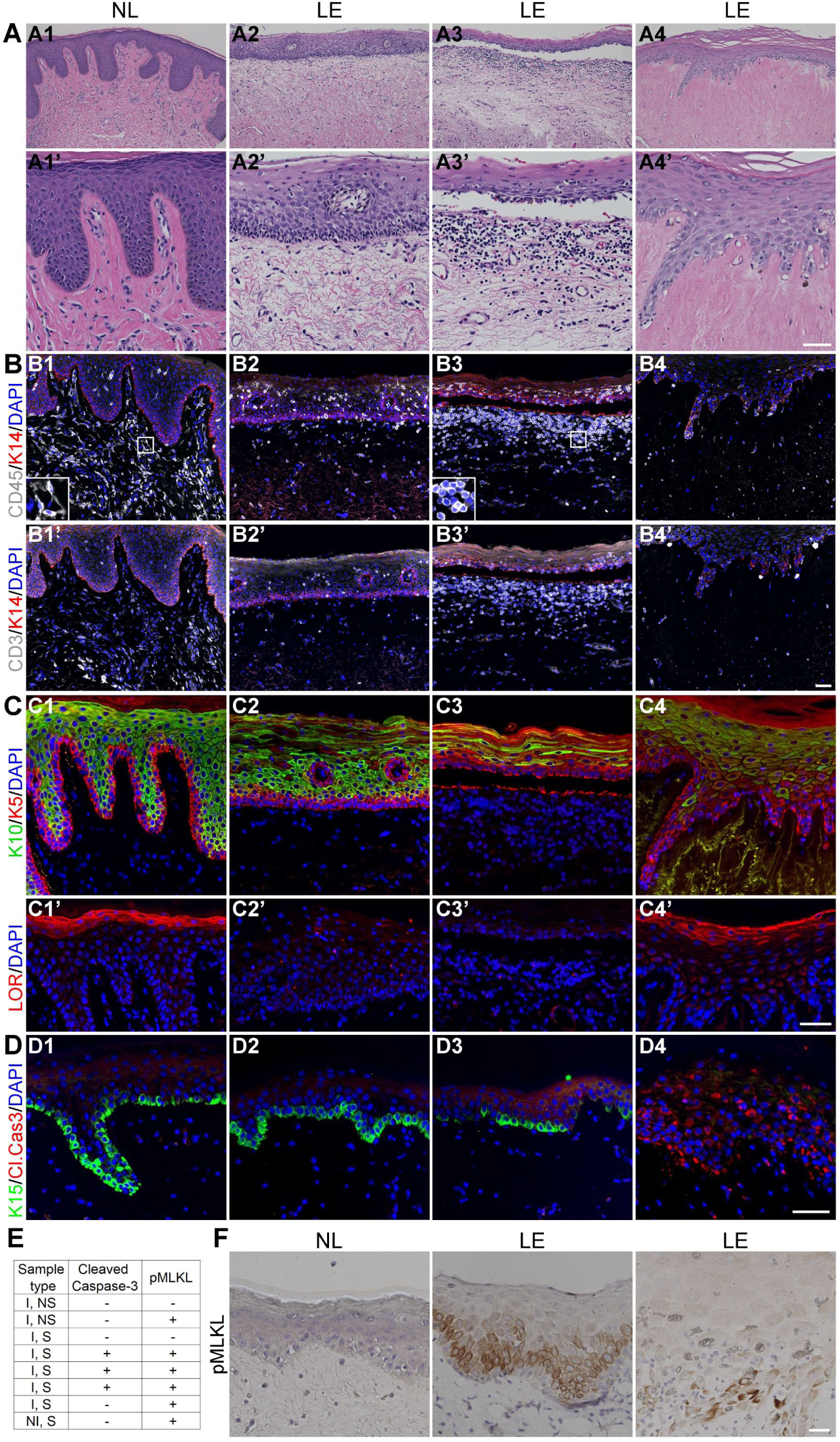
Histological hallmarks, immune infiltration, epidermal changes, and cell death in LE skin of VLS patients. (A) Representative H&E staining images of LE skin from VLS patients (A2-A4, A2’-A4’), with various levels of inflammation and sclerosis compared to NL (A1, A1’) vulvar skin. A2 and A3 show atrophic epidermis and loss of rete ridges, with dermal edema and lichenoid infiltrate. A4 shows a fully sclerotic lesion with dermal collagen homogenization. Scale bar = 160 µm in top panels, and 50 µm in bottom panels. (B-D) Immunofluorescence of NL and LE skin with the indicated antibodies. DAPI stains the nuclei. Insets in B1 and B3 show higher magnification of the boxed areas. Scale bar = 50 µm in B and C, 25 µm in D, and 16 µm for insets. (E) Summary of cell death analysis for D and F, with (F) showing representative immunohistochemical images for phosphorylated MLKL (pMLKL) in NL and LE samples. Samples were categorized per histological changes: I, inflammatory; S, sclerotic; NI, non-inflammatory; NS, non-sclerotic. Scale bar = 25 μm.

Histologically, LE samples displayed epidermal abnormalities, including pronounced hyperkeratosis, columns of parakeratosis, hypergranulosis, acantholysis of keratinocytes adjacent to the basal layer, and effacement of the epidermis (loss of rete ridges) (Fig. 1A; Table S1). Epidermal findings usually showed atrophy, although some specimens exhibited acanthosis. Additionally, there were alterations in DEJ with a lichenoid interface pattern or a subepidermal separation. Immunostaining with keratinocyte differentiation markers, K5/K14/K15 (basal), K10 (spinous), and loricrin (granular), revealed variable changes in staining pattern in the LE skin (Fig. 1C, D). Nearly all samples examined showed comparable levels of K14 expression between NL and LE skin, except for LE skin from a single patient which exhibited reduced K14 signals in basal cells but elevated staining at the skin surface (Fig. 1C; S1D). Epidermal-dermal separation in some LE samples appeared to result from basal cell lysis (Fig. 1C3). A patchy pattern of K10 staining and/or persistent K14 signals were observed in the suprabasal compartment of some LE samples (Fig. 1C). Loricrin expression varied considerably, with some LE samples showing strong and diffuse staining, while others displaying an apparent absence of expression (Fig. 1C). The presence of K15, a keratin protein typically enriched in epidermal basal stem cells^23^, was reduced or lost in six of the eight LE samples examined (Fig. 1D). This finding suggests the possibility of basal stem cell reduction in VLS.

Ki67 immunostaining did not indicate a significant reduction in epidermal proliferation in LE skin, despite the presence of epidermal atrophy (Fig. S1E). We also examined epithelial cell-adhesion marker E-cadherin and observed breakage in membrane staining in some suprabasal cells of the LE skin (Fig. S1E, F). A subset of the LE suprabasal cells lacked apparent DAPI staining, suggesting loss of nuclei (Fig. S1F). To ask if this might be due to cell death, we immunostained NL and LE skin for cleaved caspase 3, a marker of apoptosis^24^, and for phosphorylated MLKL (pMLKL), a marker of necroptosis^25^. Apoptotic cells were detected in three of the eight LE samples examined, with staining observed in basal and suprabasal keratinocytes, but not in NL skin (Fig. 1D, E). Necroptotic cells were detected in five of the eight LE samples, predominantly affecting basal keratinocytes and the suprabasal cells immediately above, but not in NL controls (Fig. 1E, F). Although necroptosis and apoptosis have traditionally been viewed as being mutually exclusive at a molecular level, they co-existed in some LE samples, though not necessarily in the same cells, and were observed in both inflammatory and sclerotic subtypes (Fig. 1D-F).

Collectively, these findings corroborate previous reports of heterogeneous changes in immune infiltration and in the dermal and epidermal compartments of VLS skin. Moreover, they underscore aberrant differentiation, loss of spatial and cell type identity, and both apoptotic and necroptotic cell death, as epidermal-associated abnormalities in VLS samples.

### GeoMx spatial transcriptomics reveals notable epidermal changes in VLS skin, highlighting a stress response

To elucidate gene expression differences in epidermal and dermal compartments of VLS LE and control skin, we analyzed clinically archived FFPE tissue specimens using a spatial transcriptomics platform, GeoMx Digital Spatial Profiler (NanoString). This platform allows for gene expression analysis of multiple regions of interest (ROIs), ranging from 13-272,074 µm^2^ in size, in a single FFPE sample^26^. We performed two independent runs, totaling seven FFPE samples: two LE and one NL from a VLS patient, one LE and NL from a second VLS patient, an unmatched LE sample from a third VLS patient, and an NL obtained from benign excision. We used fluorescent markers for pan-cytokeratin and SMA to select the ROIs, with 151 ROIs yielding quality sequencing information (Fig. 2A). These ROIs fell within seven major spatial compartments divided into epidermal and dermal regions: basal (both inter-ridge and rete ridge), inter-ridge suprabasal, rete-ridge suprabasal, papillary dermis (further segmented into SMA^+^ and SMA^-^), and reticular dermis (SMA^+^ and SMA^-^).

**Figure 2.**
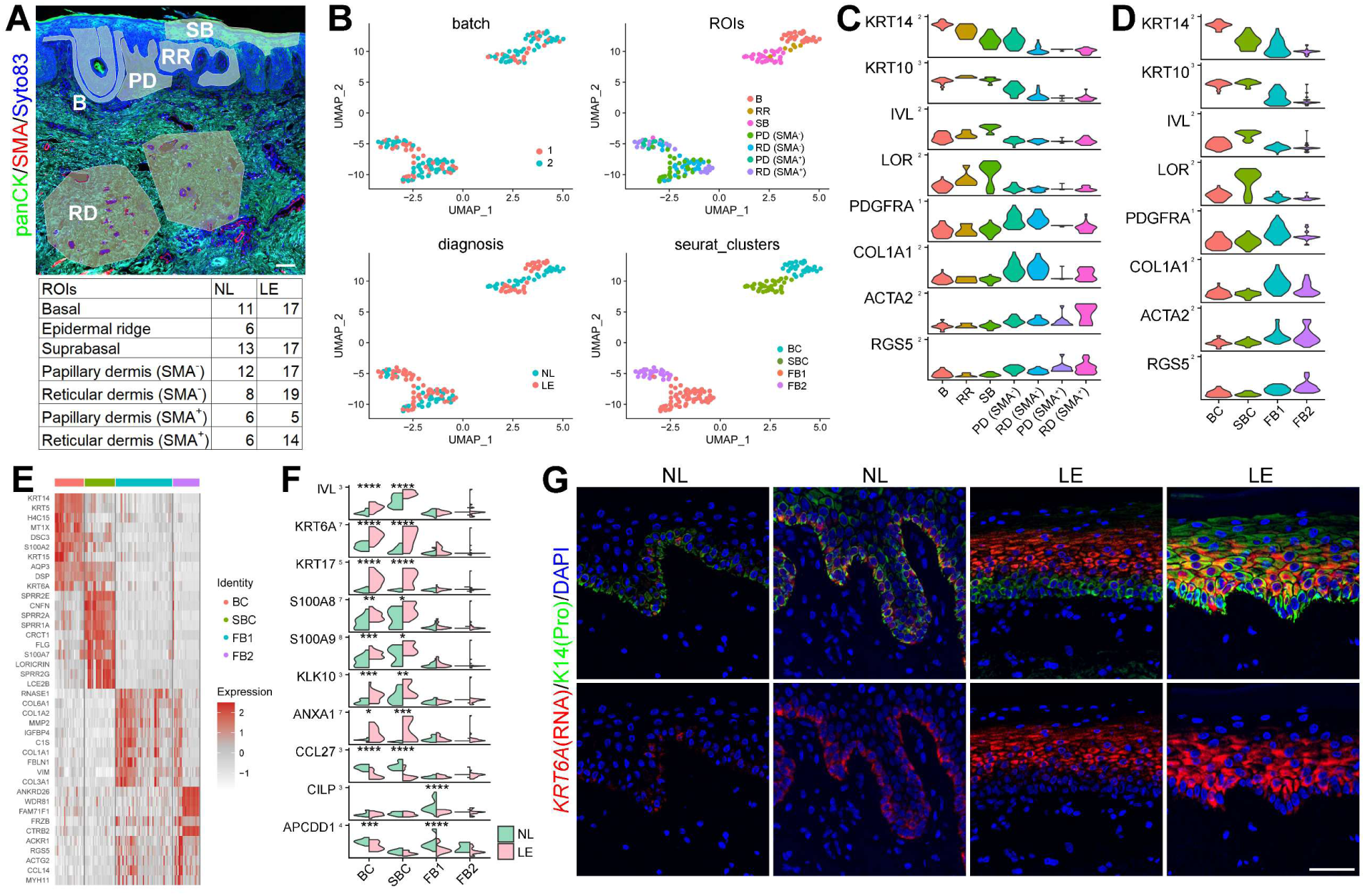
GeoMx analysis reveals gene expression differences between NL and LE skin in epidermal and dermal compartments. (A) Representative image showing ROI selection (top) and number of ROSs for each compartment (bottom). B, basal; RR, rete-ridge basal; SBC, suprabasal; PD, papillary dermis; RD, reticular dermis. panCK and SMA were used as morphology markers, and Syto83 stains the nuclei. Scale bar = 100 µm. (B) UMAPs by batch (2 independent runs), epidermal or dermal ROIs, diagnosis (NL vs. LE), or by unsupervised clustering (bottom right panel). (C-D) Violin plots showing expression of select marker genes in different ROIs (C) or Seurat clusters (D: BC, basal cells; SBC, suprabasal cells; FB1, fibroblast population 1; FB2, fibroblast population 2). (E) Heatmap of top ten marker genes for each Seurat cluster. (F) Split violin plot showing select top genes differentially expressed between NL and LE skin in each Seurat cluster. * *p* < 0.05, ** *p* < 0.01, *** *p* < 0.005, **** *p* < 0.001. (G) RNAScope data showing expanded expression of *KRT6* in LE skin. K14 protein staining marks predominantly the basal layer. Scale bar = 50 µm.

Transcriptome profiles from LE and NL skin were integrated, and uniform manifold approximation and projection (UMAP) analysis of all ROIs revealed two major clusters based on epidermal or dermal gene expression (Fig. 2B). No obvious batch effect was observed. Top markers of the spatial regions were consistent with their identity designation (Fig. 2C). Within the epidermal cluster, the rete-ridge suprabasal cells predominantly occupied intermediate positions between the basal and suprabasal cells, which were segregated into two opposite ends of the cluster (Fig. 2B). This finding suggests that the rete-ridge suprabasal cells, frequently lost in LE skin, represent an intermediate state in basal-to-suprabasal transition. Within the dermal cluster, the papillary and reticular dermal cells were not clearly segregated from each other, but a subset of the SMA^+^ dermal cells seemed to subcluster away from the rest of the dermal cells (Fig. 2B, top right).

Using Seurat for UMAP analysis, we identified four major cell clusters, two epidermal and two dermal (Fig. 2B, bottom right). The two epidermal clusters were designated basal and suprabasal using known cell-type makers (Fig. 2D, E; Table S2). Dermal clusters were designated FB1 and FB2, with FB1 being more enriched with pan-fibroblast genes (*PDGFRA*, *COL1A1*) and FB2 more with SMA^+^ cell-associated genes (*ACTA2*, *RGS5*) (Fig. 2D, E). FB1 and FB2 also showed a difference in keratinocyte-derived transcripts, possibly due to different levels of contamination with keratinocyte mRNAs. Major epidermal differences between LE and NL skin included an increased expression of terminal-differentiation gene *IVL*, stress-associated keratin genes (*KRT6A, KRT16, KRT17)*^27,28^, tissue injury-associated alarmins (*S100A8, S100A9*)^29^, *KLK10* (encoding a serine protease that is dysregulated in psoriasis)^30^, and *ANXA1* (annexin A1), as well as a decreased expression of *CCL27* (encoding a chemokine known to be decreased in multiple inflammatory skin diseases)^31^ in LE skin (Fig. 2F; Table S2). Molecular alterations in the fibroblast-enriched dermal populations were less pronounced, possibly due to greater cellular heterogeneity. *CILP*, which has been shown to antagonize the effects of TGF-β1 and IGF1^32^, as well as *APCDD1*, a Wnt signaling inhibitor^33^, were both decreased in LE skin (Fig. 2F). Interestingly, immunoglobulin (IgG) gene expression was seen in the dermis of some LE samples but not in control counterparts (Fig. S2A), suggesting that B cells might be transiently present in the LE skin.

As a proof-of-principle validation of the GeoMx-generated findings, we performed RNAScope on FFPE samples, including some that were also used for GeoMx. The results confirmed that stress keratin genes *KRT6* and *KRT16*, while expressed in the basal layer of the control skin epidermis, showed elevated or expanded expression in VLS skin, albeit to varying extents (Fig. 2G; S2B). Collectively, our data demonstrate the utility of using GeoMx on clinically achieved FFPE samples to detect skin compartment- and cell type-specific gene expression differences in LE and control skin. More importantly, these findings provide evidence for tissue injury and keratinocyte stress response in VLS skin lesions.

### scRNA-seq and single-cell spatial analyses reveal fibroblast alterations in LE skin of VLS, consistent with a pro-inflammatory and degenerative phenotype

To systemically evaluate the cellular and molecular alterations in VLS at a single-cell level, we performed scRNA-seq on patient-matched LE and NL vulvar skin samples collected from punch biopsies of five patients with biopsy-proven VLS (Table S3), using the 10x Genomics Chromium platform (Fig. 3A). One pair of samples was excluded from analysis due to low cell viability in the LE sample. Notably, this subject was the oldest and had the longest self-reported disease history, with corresponding histology showing a sclerotic phenotype - Fig. S3A; Table S3). Sequencing metrics, including the number of captured cells per biopsy sample and average reads per cell are summarized in Fig. S3B. After integrating the remaining eight samples using Seurat, correcting for batch effects, filtering low-quality cells, and removing cell doublets (see Materials and Methods), we obtained sequences from a total of 50,545 cells, with 26,938 cells from NL skin (NL1: 7,385, NL2: 12,190, NL3: 2,893, NL4: 4,470) and 23,607 cells from LE skin (LE1: 5,744, LE2: 9,452, LE3: 1,606, LE4: 6,805). UMAP clustering revealed six major cell clusters based on top expressed genes and known cell-type makers^34,35^: keratinocytes (e.g., *KRT5*, *KRT1*), immune cells (*PTPRC*), fibroblasts (*PDGFRA*), smooth muscle (SM)/pericytes (PC) (*ACTA2*), endothelial cells (*CLDN5*), and melanocytes (*PMEL*) (Fig. 3B-D; Table S3). All major cell types present in NL samples were also detected in LE samples, though with differing relative abundances (Fig. 3B; S3C).

**Figure 3.**
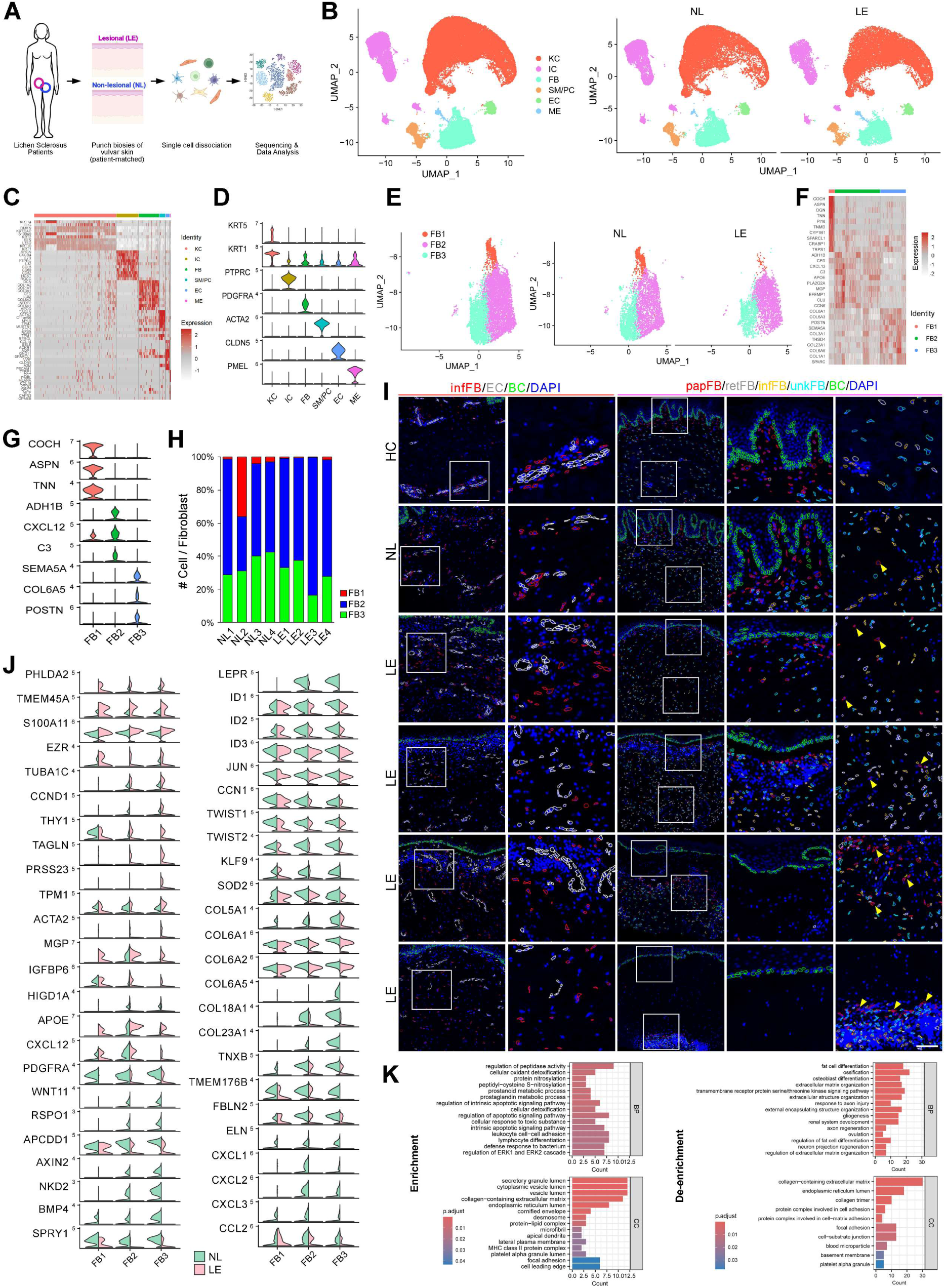
scRNA-seq workflow and overall comparison between NL and LE skin, with subclustering of fibroblasts revealing distinct subclusters and molecular differences. (A) Schematic diagram detailing the scRNA-seq workflow. Created with BioRender.com. (B) UMAP of distinct cell clusters in data integrated from all eight samples (4 NL, 4 LE) (left), and separated by NL and LE (right). KC, keratinocytes; IC, immune cells, FB, fibroblasts, SM, smooth muscle cells; PC, pericytes; EC, endothelial cells; ME, melanocytes. (C) Heatmap with top ten marker genes for each major cell type. (D) Violin plots of select markers for each major cell type. (E) UMAP of subtypes of fibroblasts in integrated data (left) and separated into NL and LE (right). (F) Heatmap with top ten marker genes for each subtype of fibroblasts. (G) Violin plot of select markers for each subtype of fibroblasts. (H) Bar plot showing relative proportions of fibroblast subclusters for all patient samples (NL and LE). (I) Xenium data revealing spatial localizations of 1) inflammatory fibroblasts (infFB) and endothelial cells (EC) on the left; and 2) four fibroblast subsets: papillary (papFB), reticular (retFB), inflammatory (infFB), and unknown (unkFB) on the right. Basal cells (BC) in epidermis are labeled green. Columns 2 and 4/5 are magnified images of insets in columns 1 and 3, respectively. Note the progressive movement of papillary fibroblasts (e.g., arrowheads in column 5) deeper in the dermis in some LE samples. Scale bar = 147 µm in column 1; 53 µm in column 2; 200 µm in column 3; 66 µm in columns 4 and 5. (J) Split violin plots of select DEGs between NL and LE skin in fibroblast subsets. (K) Diagram showing GO biological process terms enriched (left) and de-enriched (right) in LE fibroblasts compared to NL fibroblasts.

The frequent histological observation of dermal changes in VLS^7^ prompted us to first examine fibroblast changes. We computationally isolated fibroblast clusters from the integrated dataset in Fig. 3B, yielding 7,612 cells for downstream analysis (NL: 4,369, LE: 3,243). Subclustering revealed three fibroblast subpopulations: FB1 (mesenchymal), FB2 (pro-inflammatory), and FB3 (secretory), identified based on top markers for each subpopulation and markers utilized in previous studies^36^ (Fig. 3E-G). The FB3 cluster exhibited the highest expression of known papillary dermal fibroblast markers, such as *RSPO1, APCDD1, AXIN2, COL6A5, COL18A1*, and *COL23A* ^37,38^ (Fig. 3F, G; Table S4). Variation in the relative frequencies of these fibroblast subpopulations was observed in both NL and LE groups, suggesting potential influences from inter-patient variability and heterogeneity in disease onset or progression (Fig. 3H). Notably, there was a trend toward an increased proportion of the pro-inflammatory (FB2) subpopulation in LE samples (Fig. 3H). To verify that conclusions derived from the integrated master dataset held true for individual pairs of NL and LE samples before batch correction, we aggregated each NL-LE pair into separate datasets and conducted subclustering analyses on fibroblasts (Fig. S3D, E). Despite notable inter-patient variability, the pairwise comparison supported the relative increase of FB2 subpopulation and decrease in FB3 subpopulation in LE samples (Fig. S3F).

To investigate the spatial distribution of pro-inflammatory (FB2) fibroblasts, we performed Xenium single-cell spatial transcriptomics on nine FFPE LE samples and one NL sample from VLS patients, as well as one HC FFPE sample. We used Xenium’s premade skin panel (260 genes) along with 100 custom gene probes, including company-assigned markers for different fibroblast subsets (e.g., papillary, reticular, and pro-inflammatory) (Fig. S3G). A comprehensive analysis of the Xenium data will be reported separately; however, examples of individual cell type/subtypes and genes are included here. Our findings reveal that pro-inflammatory fibroblasts were predominantly located near blood vessels in the dermis of HC skin but were seen not only in vessel-rich regions but also scattered throughout the dermis in LE skin, with NL skin exhibiting an intermediate phenotype between HC and LE (Fig. 3I, left). Remarkably, while papillary fibroblasts were physically adjacent to the epidermis in HC and NL skin, they were found at progressively deeper location in the dermis of LE skin, with the fully sclerotic LE sample containing these cells (along with other fibroblast subsets) exclusively in the cellular aggregates deep in the reticular dermis (Fig. 3I, right). These observations suggest potential differentiation of perivascular fibroblasts, and/or conversion of non-perivascular fibroblasts, into pro-inflammatory fibroblasts in LE skin. Furthermore, they provide the first picture of the unusual dynamics of papillary dermal fibroblasts across different LE samples, offering mechanistic insights into dermal sclerosis in VLS.

Next, using the integrated scRNA-seq dataset, we identified differentially expressed genes (DEGs) between NL and LE skin within each fibroblast subsets (Table S4). Compared to other cell types in LE skin, fibroblasts exhibited particularly high levels of transcripts typically restricted to keratinocytes (e.g., *KRT14*), likely due to contamination with keratinocyte-derived ambient RNAs. These keratinocyte transcripts (Table S4) were manually removed from subsequent analysis.

Importantly, LE fibroblasts showed broad upregulation of genes associated with cellular stress and death, such as *PHLDA2* (a mediator of ferroptosis)^39^, *TMEM45A* (a context-dependent apoptosis regulator)^40^, *S100A11* (an alarmin that can induce chemokine response through engaging its RAGE receptor)^41^, *EZR* (a cytoskeleton-associated protein involved in cell shape-sensing and hypoxia-induced autophagy pathways)^42^, and *TUBA1C* (a PANoptosis-related predictor in glioma)^43^ (Fig. 3J). Additionally, genes with roles in cell cycle progression (*CCND1*) or in regulation of fibrosis and extracellular matrix (ECM) remodeling, including *THY1*, *TAGLN*, *PRSS23*, *TPM1*, *ACTA2*, *MGP*, and *IGFBP6*^44–50^, were increased in LE (Fig. 3J). Among these, *THY1*, *TAGLN*, *TPM1*, and *ACTA2* are known markers of myofibroblasts^51,52^, suggesting an “activated” state of LE fibroblasts.

*APOE*, previously reported to be expressed in pro-inflammatory fibroblasts^35^, was enriched in FB2 subpopulation and showed increased expression across all fibroblast subsets in LE samples (Fig. 3J). Similarly, *CXCL12*, an FB2-enriched cytokine, was increased in all LE fibroblast subsets (Fig. 3J). These findings corroborate the Xenium results to suggest the acquisition of a pro-inflammatory state by non-inflammatory fibroblasts in diseased skin.

Overall, there were more DEGs with significantly decreased expression in LE fibroblasts than those with increased expression (Table S4). Particularly notable was the decreased expression of developmental signaling genes including *PDGFRA*, which plays a role in fibroblast proliferation and fibroblast-to-myofibroblast transition^53^, Wnt agonists (*WNT11, RSPO1*) and antagonists (*APCDD1, AXIN2, NKD2*), *BMP4*, *SPRY1,* and *LEPR* (Fig. 3J). Decreased expression was also seen for several mitogen-induced early response genes (i.e., immediate early genes) such as *ID1, ID2, ID3, JUN,* and *CCN1*^54,55^, EMT-promoting transcription factors *TWIST1/2*^56^, oxidative stress-associated genes *KLF9* and *SOD2*^57^ (Fig. 3J). Multiple structural genes were decreased in LE, including collagen genes *COL5A1, COL6A1/2, COL6A5, COL18A1, COL23A1; TNXB* and *TMEM176B* involved in ECM organization; *FBLN2,* an ECM-secreted protein involved in basement membrane zone stability^58^; and *ELN* that encodes elastin (Fig. 3J). Intriguingly, a number of these downregulated genes are normally enriched in papillary fibroblasts. Moreover, the expression of several neutrophil- or monocyte-attracting chemokines, *CXCL1/2/3* and *CCL2*, was decreased in LE fibroblasts (Fig. 3J), correlating with the apparent scarcity of neutrophils and monocytes in LE (see below). We also investigated the expression of the key DEGs using individually paired NL and LE aggregations by patient, confirming the consistency of our overall findings (Fig. S3H).

To further assess changes in fibroblast function, we conducted gene ontology (GO) analysis. The analysis revealed an enrichment in terms including “regulation of protease activity”, “cellular oxidant detoxification”, and “regulation of apoptotic pathway”, as well as a de-enrichment of terms including “fat cell differentiation”, “extracellular matrix organization”, and “collagen-containing extracellular matrix”, in LE skin (Fig. 3K). Overall, the gene expression changes observed in LE fibroblasts indicate the acquisition of a pro-inflammatory and degenerative phenotype, at the apparent expense of the developmental, differentiation, and secretory roles of NL fibroblasts. Given the potential for fibroblasts to differentiate into adipogenic precursors^59^, these findings may also indicate the loss of a fibroblast progenitor state in LE skin. Moreover, the increased expression of some inflammatory factors such as *CXCL12,* coupled with reduced expression of others like *CXCL1/2/3,* suggests that fibroblasts may contribute to instructing a chronic inflammatory state rather than an acute one in stable VLS.

### Immune alterations in VLS are complex and predominantly T cell-driven, suggesting a preexisting blueprint in NL skin, with a cytotoxic and activated phenotype in LE skin

Immune-mediated mechanisms, particularly those associated with a Th1-specific IFN-γ-induced phenotype, have been implicated in VLS pathogenesis^7^. To evaluate immune alterations in VLS, we computationally isolated all immune cell clusters from the integrated dataset, comprising 8,314 total cells (NL: 5,368, LE: 2,946). Downstream subclustering analysis yielded a total of seven subpopulations: CD4^+^ T1, CD4^+^ T2, CD8^+^ T1, CD8^+^ T2 (potentially including natural killer T cells), cycling T cells (cycTC), dendritic cells (DC)/ macrophages (MP), and mast cells (MC) (Fig. S4A-C). Variations in the relative frequency of the cell subpopulations were seen in both NL and LE skin, with T cells being the most prominent cell types (>75%) in all samples (Fig. S4D). When individual pairs of NL and LE data were aggregated, clustering analysis revealed the presence of additional immune cell types such as B cells (pair 2) and natural killer cells (pair 4), which were not consistently observed across all samples (Fig. S4E-G). In both integrative and pair-wise analyses, we did not observe a consistent expansion or reduction of any given immune cell type from NL to LE (Fig. S4D, G). These data suggest that an LE-like immune blueprint preexists in NL skin and the immune response in LE skin is heterogeneous and spatiotemporally dynamic. These “snapshots” may indicate the dynamic nature of the disease, with varying phases of inflammation and T cell activity.

Next, we identified DEGs between NL and LE skin in each of the immune cell clusters (Fig. S4H; Table S5). Of interest, *NEAT1*, a long-coding RNA that facilitates inflammasome activation in macrophages, modulates T cell differentiation, and promotes Th2 cytokine production in CD4^+^ T cells^60–62^, exhibited increased expression in all T cell subsets and DC/MPs of LE compared to NL (Fig. S4H). *DUSP4*, a negative regulator of MAPK pathways in T cells^63^, demonstrated elevated expression in most T cell subsets and DC/MP cells in LE compared to NL (Fig. S4H). Importantly, *CD69*, historically thought as an early activation marker of T cells and other immune cells and recently recognized as a canonical marker of tissue-resident memory T cells^64–66^, demonstrated consistent downregulation across nearly all LE immune cell populations (Fig. S4H). Reduced expression of *PHLDA1, HSPA1A, HSPA1B, FOS, CITED2, DUSP1*, and *TUBA1A* - genes with relevance to cellular stress and early response^15^, was observed in CD4^+^ and CD8^+^ T cells across most if not all LE samples (Fig. S4H).

DEGs with more localized changes, identified in particular immune cell subsets, were also noted. *TNFRSF4*, normally expressed in activated CD4^+^ and CD8 ^+^ T cells^67^, was upregulated in CD4^+^ T2 cells in most LE samples (Fig. S4H). In general, CD8^+^ T cells exhibited more consistently altered DEGs between LE and NL samples compared to CD4^+^ T cells (Fig. S4H). CD8^+^ T1 cells of all LE samples showed elevated expression of *PDE4D, HLA-DRB1, RGS1*, and *PTMS*, whereas their expression of genes such as *NKG7, CD74,* and *RNF213* was upregulated in most LE samples (Fig. S4H). *PDE4D* activity is recognized to enhance the activation and function of CD8^+^ T cells^68^, and *RGS1* and *PTMS* have been implicated in dysregulated immune responses^69^. *HLA-DRB1* encodes a major histocompatibility complex (MHC) class II gene, and its elevated expression in LE CD8^+^ T1 cells suggests that these cells may participate in antigen presentation to CD4^+^ T cells.

Remarkably, CD8^+^ T2 cells, characterized by high expression of cytotoxicity genes such as *GZMB*, were already detected in all NL samples (Fig. S4D, H). However, *GZMB* levels in CD8^+^ T2 cells were higher in LE compared to NL counterparts in most VLS patients examined (Fig. S4H). Surprisingly, decreased expression of *TNF* and *IFNG* (IFN-γ), encoding two classical Th1 inflammatory factors important in CD8^+^ T cell function, was observed in CD8^+^ T2 cells from most LE samples (Fig. S4H). CD8^+^ T1 cells in LE samples also showed consistently decreased expression of genes encoding inflammatory factors such as *LTB, AREG,* and *TNF* (Fig. S4H). To further understand these findings, we examined the expression of known markers of T cell exhaustion and chronic activation^70,71^. The expression of *LAG3*, *ITGAE*, *CTLA4*, and *HAVCR2* was elevated in CD8^+^ T2 cells in at least three of the four LE samples (Fig. S4I). Together, these data suggest that the cytotoxic CD8 ^+^ T cells in some LE skin are likely in a prolonged activation state.

The DC/MP cluster represents a small population and subclustering attempts failed to resolve discrete subsets. However, reduced expression of classic DC markers *CD1C* and *CD1B* was observed in most LE samples compared to NL (Fig. S4H), consistent with previous report of reduced DC frequency in LS skin dermis^72^. Moreover, the DC/MP cluster in all LE samples showed reduced expression of *IL1R2,* which encodes a decoy receptor for the inflammatory cytokine IL-1^73–75^ (Fig. S4H), suggesting elevated IL-1 signaling in LE skin. We also observed MC-specific changes, with *SFN, CSTA, DMKN, HLA-B,* and *HLA-C* showing elevated expression across all LE samples (Fig. S4H). Given that IFN-γ–primed skin mast cells can act as antigen-presenting cells^19,20^, the increased MHC class I gene (*HLA-B/C*) expression suggests that mast cells may contribute to antigen presentation to CD8^+^ T cells in LE skin.

GO analysis of all immune cells revealed enrichment of terms such as “leukocyte cell-cell adhesion” and “antigen processing and presentation via MHC class II”, as well as de-enrichment of terms such as “response to corticosteroid”, “regulation of DNA-binding transcription factor activity”, and “activation of immune response” in LE skin (Fig. S4J). Together, our findings underscore the coexistence of elevated adaptive immune response and aberrant inflammation in VLS, suggesting that by the time lesions appear, immune activities are already advanced. Increased cytotoxicity and *LAG3* expression in CD8^+^ T cells of LE skin indicate a highly activated T cell phenotype. Additionally, CD69 expression in CD8^+^ T cells of NL skin suggests that these cells might already be poised to respond to antigens and induce LE pathology.

### Keratinocyte and melanocyte alterations in LE epidermis suggest epidermal stem/progenitor cell depletion, premature terminal differentiation, stress response, and elevated interferon (IFN) signaling

Keratinocytes, the predominant cell type in our analysis, have been implicated in immune-mediated disease, serving both as sources and targets of inflammatory mediators^76^. To investigate keratinocyte differences between LE and NL skin, we isolated all keratinocyte clusters from the integrated dataset, resulting in a total of 30,604 keratinocytes: 15,061 from NL skin and 15,543 from LE skin. We identified six distinct keratinocyte subsets according to their top markers: three basal subtypes – basal stem cells (BSC), cycling basal cells (cycBC), and bulk basal cells (BC) – as well as three differentiating subtypes – spinous cells (SC) 1 and 2, and granular cells (GC) (Fig. 4A-C). This organization in the vulvar skin is consistent with epidermal compartments observed in human skin from other body regions, such as foreskin and hip skin^77,78^, but with distinctions (Fig. S5A). In LE skin, there was a notable increase in the more differentiated keratinocyte subtypes, specifically the SC2 and GC subpopulations (average for NL: 19.3%, LE: 42.8% for SC2; NL: 7.9%, LE: 16.5% for GC) (Fig. 4D; S5A). This was accompanied by a decrease in the basal and early spinous subpopulations (Fig. 4D; S5A).

**Figure 4.**
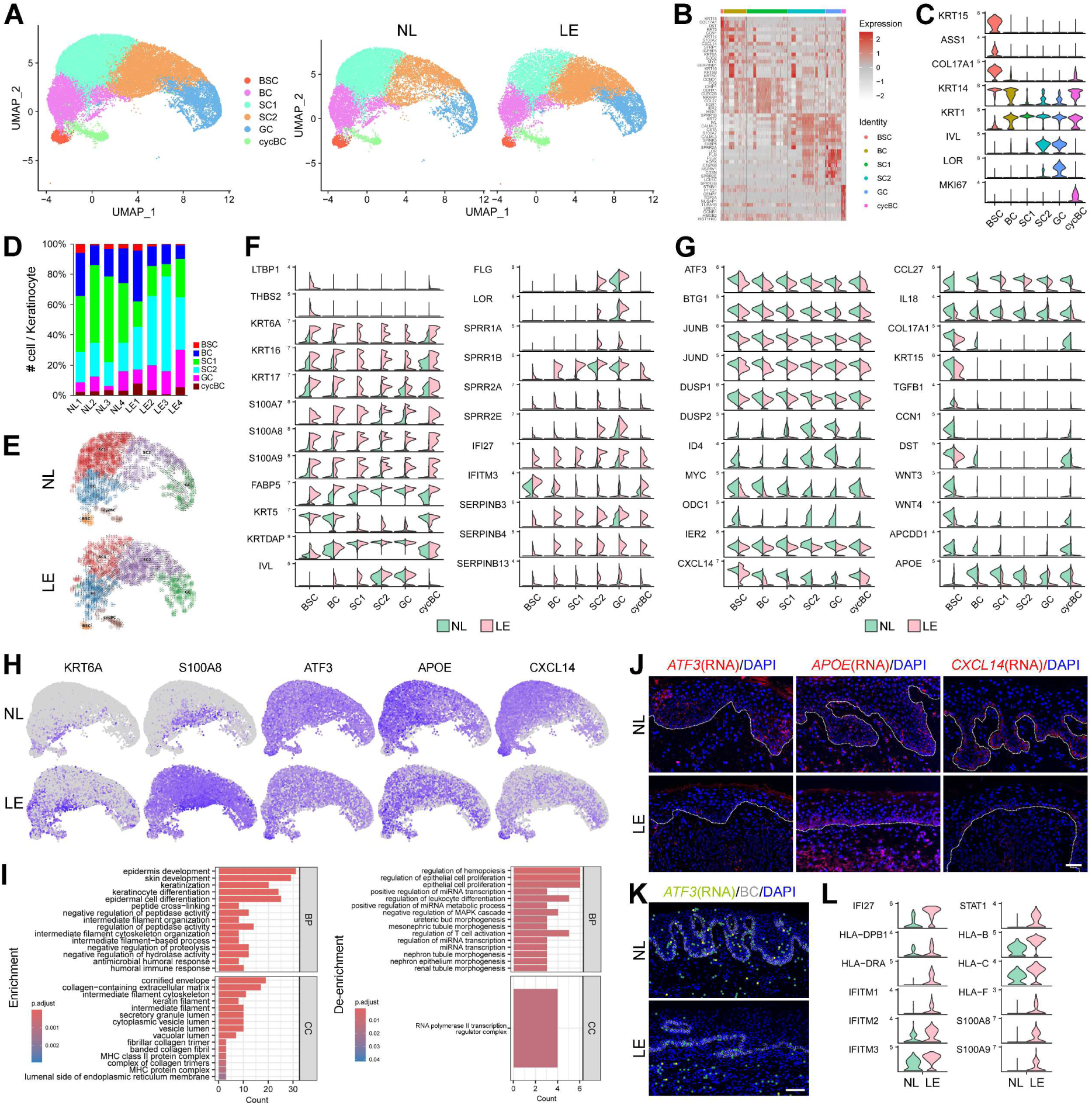
Molecular alterations in LE keratinocytes and melanocytes. (A) UMAP of subtypes of keratinocytes in integrated data (left) and separated into NL and LE (right). BSC, basal stem cells; BC, basal cells; spinous cells (SC); granular cells (GC); and cycling basal cells (cycBC). (B) Heatmap with top ten marker genes for each subtype of keratinocytes. (C) Violin plot of select markers for each subtype of keratinocytes. (D) Bar plot showing relative proportions of keratinocyte subclusters for all patient samples. (E) Projection of TFvelo velocity fields onto the UMAP space of keratinocytes in NL and LE. (F-G) Split violin plots of select genes upregulated (F) or downregulated (G) in LE keratinocyte subsets compared to NL counterparts. (H) Feature plots of the indicated genes in NL and LE keratinocytes. (I) Diagram showing GO biological process terms enriched (left) and de-enriched (right) in in LE keratinocytes compared to NL keratinocytes. (J) RNAScope data on *ATF3*, *CXCL14*, and *APOE* transcripts in NL and LE skin. Dotted line indicates basement membrane. Scale bar = 50 µm. (K) Density plot with spatial localization of *ATF3* expression in NL and LE skin by Xenium analysis. Yellow signifies higher expression and green marks lower expression levels. BC, basal cells. Scale bar = 100 µm. (L) Violin plots of select DEGs between NL and LE melanocytes.

To further investigate the differentiation abnormalities in LE skin, we employed RNA velocity, a computational method that predicts the future state of individual cells by assessing the relative abundance of spliced and unspliced transcripts in single-cell data^79^. Velocity calculations using both linear and non-linear methods revealed major differences in the differentiation trajectories of keratinocytes between NL and LE samples (Fig. S5B). Both models identified an absence of “backward” transitions from SC to BC, and indicated a relative “inertness” of SC2 and GC populations in LE skin compared to NL (Fig. S5B). Additionally, we employed a modified velocity analysis method, known as TFvelo^80^, which uses scRNA-seq data based on transcription factor (TF)-target relation and incorporates prior knowledge from a TF-target database^81^. This method provided a more robust projection of epidermal differentiation trajectories, while corroborating the observed differences in SC “reversibility” and GC “inactivity” between NL and LE skin (Fig. 4E). These findings suggest reduced plasticity in the early stages and altered differentiation trajectories in the later stages of keratinocyte differentiation within LE skin.

In general, analysis of the pair-wise aggregation data corroborated the decrease in basal and/or early spinous keratinocytes and the increase in late differentiating keratinocytes in LE, but with heterogeneous manifestations across patient samples as well as the emergence of aberrant differentiation “trajectories” in some LE samples (Fig. S5C-E). These data, along with the observed loss of rete ridges containing basal-suprabasal intermediate states (see GeoMx analysis above), suggest the depletion of stem/progenitor and transitional cells, along with premature terminal differentiation, as part of the epidermal phenotype in LE skin.

We next identified DEGs within each of the keratinocyte subtypes between NL and LE skin. Table S6 shows the top DEGs up- or down-regulated in at least one keratinocyte subset of LE skin compared to its NL counterpart. Some subset-specific changes were detected, such as elevated expression of *LTBP1* and *THBS2,* and reduced expression of *KRT15,* in LE BSCs (Fig. 4F, G). Importantly and corroborating the GeoMx data, we observed upregulation of tissue damage/stress-associated genes, including stress keratins *KRT6A/B/C, KRT16, KRT17*, and alarmins *S100A7*, *S100A8, S100A9*, across multiple LE keratinocyte subsets (Fig 4F, H). Expression of *FABP5* was also elevated, universally, in keratinocyte subsets (Fig. 4F). Variable changes in classical epidermal differentiation markers such as *KRT5/14, KRTDAP, IVL,* and *FLG* were noted (Fig. 4F), likely reflecting heterogeneity in epidermal fluxes due to constant damage and repair throughout the disease course. Particularly notable was the elevated expression of late differentiation genes such as *FLG, LOR* and *SPRR* family members across multiple keratinocyte subsets in multiple LE samples (Fig. 4F), supporting the notion of premature epidermal differentiation. Finally, the expression *IFI27* and *IFITM3*, two IFN-induced genes^82^, was elevated in multiple LE keratinocyte subsets (Fig. 4F). Other upregulated genes across multiple subsets of keratinocytes included *SERPINB3/4/14* (Fig. 4F), which are known to be induced in inflammatory and hypoxic states^83^.

We also identified downregulated DEGs across multiple keratinocyte subsets in LE skin compared to NL, which included several early response genes such as *ATF3*^84^, *BTG1*, *JUNB/D, DUSP1/2*^63^*, ID4*^85^*, MYC*^86^, *ODC1*, and *IER2* (Fig. 4G, H)^63,85–90^. Also, among the top downregulated genes were those encoding chemokines *CXCL14* and *CCL27,* and proinflammatory cytokine *IL18*^91^ (Fig. 4G). Some genes, such as *COL17A1*, *KRT15*, *TGFB1, CCN1, DST,* as well as Wnt signaling ligands/inhibitor *WNT3/4* and *APCDD1,* showed reduced expression predominantly in basal and differentiating keratinocytes (Fig. 4G). Moreover, *APOE*, encoding a fat-binding protein involved in cholesterol/fat metabolism and implicated in skin inflammation^92^, showed reduced expression predominantly in cycling BC and SC1 subsets (Fig. 4G, H). Finally, GO analysis of all keratinocytes revealed an enrichment of terms such as “epidermal development”, “keratinization” and “keratinocyte differentiation”, and de-enrichment of terms such as “regulation of hematopoiesis” and “epithelial cell proliferation” in LE skin compared to NL skin (Fig. 4I).

Using RNAScope, we found *ATF3* and *APOE* to be expressed in both basal and spinous cells, and *CXCL14* expressed more prominent in basal cells, of NL skin (Fig. 4J), consistent with their expression patterns along the UMAP trajectory (Fig. 4H). Importantly, expression of *ATF3* and *APOE* was decreased in the suprabasal compartment of LE skin compared to NL skin (Fig. 4J). Reduction in *CXCL14* expression in LE skin was more variable, but occurred in at least some LE samples (Fig. 4J). *ATF3* was included as one of the 100 custom probes in our Xenium analysis, and while its expression was detected in basal and, to a greater extent, suprabasal cells of HC and NL skin, all LE samples exhibited reduced *ATF3* expression in their suprabasal compartment (Fig. 4K).

VLS often presents with depigmentation or hypopigmentation and is frequently associated with vitiligo, an immune-mediated depigmenting skin condition^93^. Despite reports of melanocyte depletion^94^ and reduced transfer of melanosomes to keratinocytes in VLS^95^, studies investigating melanocyte involvement in VLS remain limited. Melanocytes typically reside in the basal layer of the epidermis^96^ and are influenced by adhesion and paracrine signaling interactions with keratinocytes^97,98^. We computationally isolated melanocytes from the integrated dataset (Fig. 3B) for detailed analysis. Despite the small number of melanocytes detected (NL: 122; LE: 233), we found increased IFN I target gene signatures (*IFI27, IFITM1, IFITM2, IFITM3, IFI6*) and increased *STAT1* expression in melanocytes of most LE samples compared to NL (Fig. 4L). We also observed increased expression of MHC class I (*HLA-B, HLA-C, HLA-F*), which are induced by IFN signaling^99^, and class II (*HLA-DPB1, HLA-DRA*) genes in LE melanocytes (Fig. 4L). These findings highlight melanocytes as a significant site of elevated IFN signaling^100^, as well as their potential involvement in antigen-presentation to both CD4^+^ and CD8^+^ T cells. In contrast, *HLA-A/B/C* genes were not consistently altered in LE keratinocytes compared to their NL counterparts (Fig. S5G).

Collectively, these findings indicate reduced expression of genes linked to epidermal stem/progenitor and transitional states, along with premature expression of terminal differentiation genes in LE skin. They also provide evidence for keratinocyte stress and alarmin production. The molecular changes in both keratinocytes and melanocytes suggest elevated IFN signaling within the epidermal compartment of LE skin and implicate melanocytes in antigen presentation.

### Analyses of ligand-receptor expression and downstream signaling predict altered cell-cell signaling communications and elevated IFN/JAK/STAT pathway activation in LE skin

To probe global differences in cell-cell communications in LE compared to NL skin, we performed comparative CellChat analysis to study the signaling interaction potential among all major cell types^101^. CellChat infers cell-cell communication networks based on the expression levels of ligands, receptors, and their cofactors. We used individual keratinocyte, immune cell, and fibroblast subsets, leading to 20 cell groups for analysis.

We first examined the total number of potential cell-cell communications and their prominently affected cell type/subtypes. Overall, there appeared to be a higher number of possible interactions in NL skin than in LE skin among or within the different cell groups (Fig. 5A). Putative signaling from all keratinocyte subpopulations to all immune cell subpopulations (except for MCs), particularly from BSC and BC to CD8^+^ T1 and cycTC, showed increased strength in LE skin (Fig. 5B). In fact, the CD8^+^ T1 and cycTC immune subpopulations were strong receivers of signals from most of the cell populations in LE skin (Fig. 5B). Elevated signaling strength from keratinocytes to melanocytes and from CD4^+^ T2 cells, smooth muscle cells, pericytes and melanocytes to melanocytes was also observed (Fig. 5B). These findings underscore BSC/BC keratinocytes as among the most activated (relative to NL) signaling senders, and CD8^+^ T1, cycTC, and melanocytes the most activated signaling receivers, in the LE skin microenvironment.

**Figure 5.**
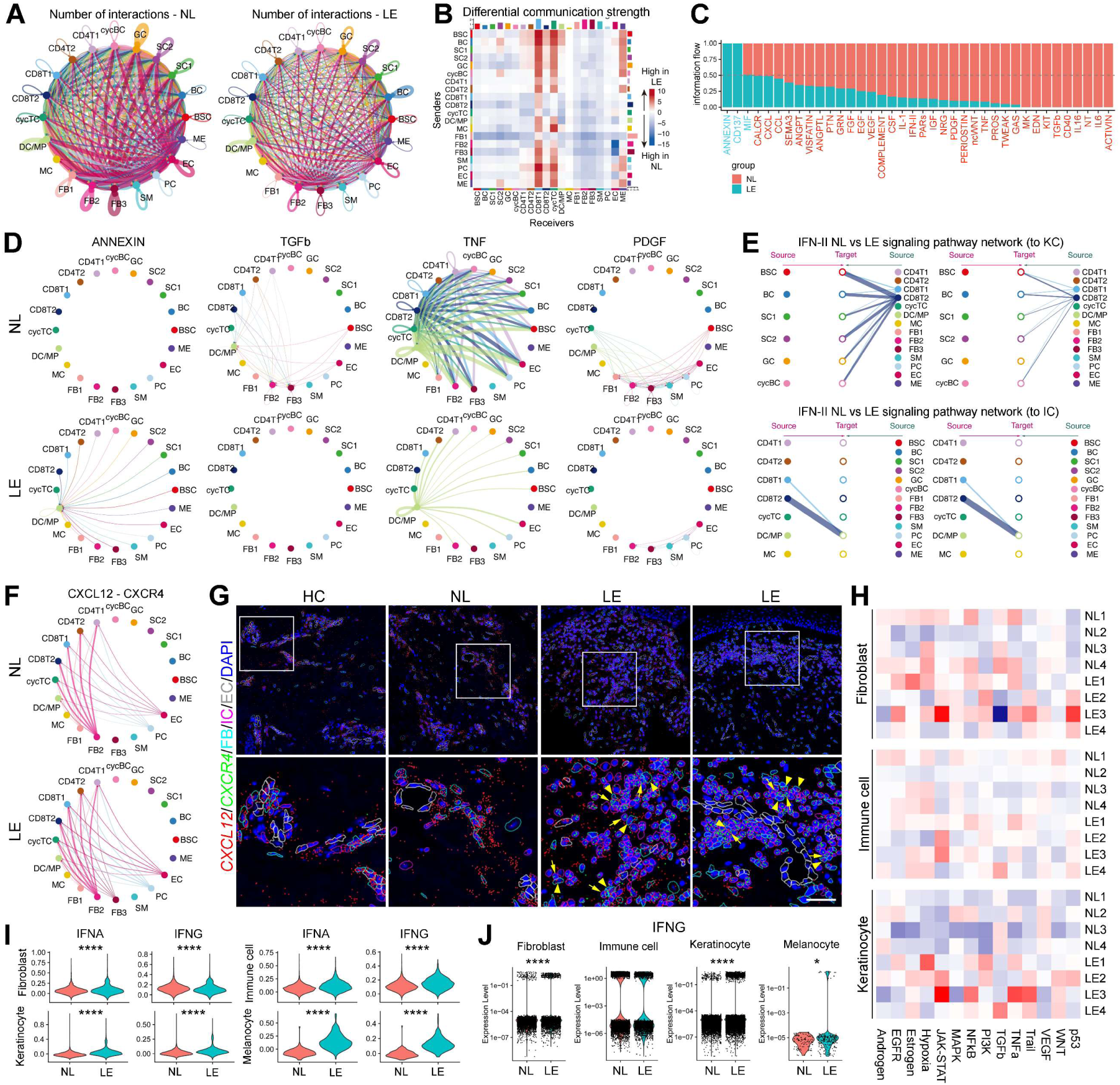
Cell-cell communication and pathway activity changes in LE skin. (A) Circle plot (by count) revealing the number of putative ligand-receptor interactions between cell groups for NL and LE; line thickness represents strength of communication. (B) Heatmap (by weight) summarizing differential strength of communication in LE compared to NL. Red (positive values) and blue (negative values) in the color bar indicate a higher number of predicted interactions in LE and NL skin, respectively. (C) Significant signaling pathways were ranked based on their differences of overall information flow, calculated by summarizing all communication probabilities in a given inferred network. Those colored red and green are more enriched in NL and LE skin, respectively. (D) Chord diagrams of select inferred signaling networks. (E) Hierarchy plots of IFN II signaling from all cells to keratinocytes (top) or to immune cells (bottom) in NL (left) vs. LE (right) skin. (F) Chord diagrams of the inferred CXCL12-CXCR4 signaling network in NL vs. LE skin. (G) Xenium data showing spatial localization of *CXCL12* and *CXCR4* transcripts with identification of cell types including fibroblasts (FB), immune cells (IC), and endothelial cells (EC). Bottom row shows high magnification images of boxed areas in the top row. Arrowheads and arrows indicate neighboring *CXCL12*- and *CXCR4*-expressing cells, respectively. Scale bar = 100 µm in top row and 33 µm in bottom row. (H) PROGENy analysis of signaling pathway activation in fibroblasts, immune cells, and keratinocytes using scRNA-seq data for all patient samples. (I) Gene scoring analysis of the indicated cell types in NL vs. LE using MSigDB signatures for HALLMARK_INTERFERON_ALPHA_RESPONSE (IFNA) and HALLMARK_INTERFERON_GAMMA_RESPONSE (IFNG). **** *p* < 0.001. (J) Violin plots of *IFNG* expression by cell type in NL compared to LE, visualized by overlaying a jitter plot and transforming data using a log10 scale. * *p* < 0.05, **** *p* < 0.001.

As LE fibroblast contamination with keratinocyte-derived transcripts may distort signaling predictions, we subclustered the NL+LE master dataset at a high resolution to enable computational removal of nearly all “contaminated” fibroblasts, and repeated CellChat analysis. The major conclusions, such as CD8^+^ T1, cycTC, and melanocytes being the most activated signaling receivers in LE skin, remained the same (Fig. S6A). Interestingly however, the inflammatory fibroblast subpopulation (FB2) now emerged as the strongest signaling senders in LE skin, with keratinocytes, immune cells, FB2 fibroblasts, and pericytes as their favored signaling receivers (Fig. S6A). Thus, FB2 fibroblasts are likely also key participants in cell-cell signaling communications in LE skin.

Next, we compared the information flow, computed as the sum of communication probabilities among all pairs of cell groups in the inferred network, for specific signaling pathways between LE and NL skin. Considerable variability was observed across patient samples. Of the 41 pathways detected by CellChat, 38 exhibited consistent changes in signaling strengths across both integrated data and at least two pair-wise comparisons (Fig. 5C). Notably, ANNEXIN and CD137 (TNFSF9) pathways showed increased information flow in LE samples compared to NL controls (Fig. 5C, D; S6B). The increased ANNEXIN signaling in LE skin is in part attributed to elevated expression of ANNEXIN 1 (*ANXA1*) in keratinocytes, along with elevated expression of *FRP1*, but not *FRP2*, receptor gene in keratinocytes (at very low frequency) and immune cells (particularly DC/MPs) (Fig. S6C). Increased CD137 signaling in LE skin involved CD8^+^ T2 and cycTC as the signal sender or receiver (Fig. S6B), consistent with CD137-CD137L being a costimulatory pathway for CD8^+^ T cells and serving as a critical immune checkpoint with implications for autoimmunity^102–104^.

Other pathways, such as TGFb, TNF, PDGF, and IFN-II, showed overall reduced information flow in LE skin (Fig. 5C-E). TGFb signaling in NL skin involved various senders and three primary receivers: FB2, FB3, and DC/MP, with information flow through all signaling routes diminished in LE skin (Fig. 5D). TNF signaling was widespread across many cell types in NL skin but became restricted in LE, with DC/MPs being the only signaling receivers (Fig. 5D). Fibroblasts, smooth muscle cells, and pericytes were the predominant signal receivers of PDGF signaling in NL skin, but the overall signaling flow to them was decreased in LE skin (Fig. 5D). Finally, while CD8^+^ T cells were the major source of IFN-II signaling in NL skin, overall signaling potential was predicted to decrease in LE skin (Fig. 5E). Collectively, these findings suggest that both signaling strength and architecture of putative cell-cell communication networks are altered in LE compared to NL skin.

Intrigued by the increased expression of *CXCL12* in LE fibroblasts, we examined its possible signaling interactions. CellChat predicted that CXCL12-CXCR4 signaling predominantly occurs from FB2 fibroblasts to T cell subsets, with potential signaling expanded and FB3 emerging as a new signaling source in LE compared to NL skin (Fig. 5F). Finer analysis revealed a slight increase in CXCL12-CXCR4 communication probability between FB2 and CD8^+^ T cells, but a decrease between FB2 and CD4^+^ T cells, in LE skin (Fig. S6D). To place these changes in a spatial context, we examined in situ *CXCL12* and *CXCR4* expression using our Xenium data. In HC and NL skin*, CXCL12* expression was predominantly localized to dermal cells surrounding blood vessels, whereas *CXCR4* expression was mainly in immune cells (Fig. 5G). In nearly all LE samples examined, *CXCL12*-expressing dermal cells were found not only around blood vessels but also dispersed among dermal fibroblasts (Fig. 5G). Furthermore, elevated frequencies of *CXCR4-*expressing immune cells were observed in six of the nine LE skin samples examined (Fig. 5G). Although *CXCL12*-expressing fibroblasts and *CXCR4*-expressing immune cells were not exclusively associated, they were often found in close proximity within LE skin, suggesting a likelihood of signaling interactions. CXCR4 is a marker for tissue-resident CD4^+^ and CD8^+^memory antigen-experienced T cells^105^, and this interaction with CXCL12-expressing fibroblasts may be a mechanism of LE pathology.

While CellChat can predict signaling potential, the inferred interactions are hypothetical, and more definitive evidence of signaling alteration requires analyzing the downstream signaling output for each pathway. Therefore, we applied PROGENy, a computational pipeline that infers pathway activity from gene expression using publicly available perturbation experiments^106^, to the integrated master dataset to analyze the activities of 14 signaling pathways. Overall, differential signaling activities between LE and NL skin were noted across fibroblast, immune cell, or keratinocyte subsets, with considerable heterogeneity observed among patients and across cell type/subtypes (Fig. 5H; S6E). Specifically, both PI3K and p53 pathways were elevated in all fibroblast subsets of all LE samples compared to NL (Fig. 5H). Conversely, androgen and TGF-b pathways were decreased in LE fibroblasts compared to NL across all samples. PI3K/AKT signaling plays a role in numerous cellular responses, including proliferation and apoptosis, and exhibits diverse crosstalk mechanisms^107^. p53 pathway is activated by a myriad of cellular stressors (e.g., DNA damage, oxidative stress) to then terminate cellular processes, such as cell cycle and differentiation, and promote apoptosis^108^. The reduced TGF-b signaling activity is unexpected, given that it is one of the most well-characterized pro-fibrosis pathways^109^. LE immune cells also exhibited elevated p53 pathway compared to their NL counterparts, along with increased JAK/STAT activity in most LE samples (Fig. 5H). In LE keratinocytes compared to their NL counterparts, the PI3K, JAK/STAT, estrogen, trail, and Wnt pathways were elevated, while the VEGF pathway was decreased (Fig. 5H). Of note, the LE sample with the least JAK/STAT activation exhibited the mildest histological abnormalities, characterized by the absence of dermal sclerosis, retention of epidermal rete ridges, and only minimal immune cell infiltration (Fig. S6F).

JAK/STAT pathway, involved in many vital cellular processes including cancer and inflammation, is linked to various autoimmune diseases and cancers when dysregulated^110^. IFNs are among the many cytokines and growth factors that can activate JAK/STAT signaling^110^. Given the activation of IFN-γ/JAK/STAT pathway in other lichenoid inflammatory disorders (e.g. lichen planus)^111^ and immunohistochemical evidence of increased IFN-γ in VLS^17^, which seems to contrast our finding of reduced IFN-γ signaling probability in LE skin, we set out to evaluate type I and type II IFN target signatures in our scRNA-seq data. MSigDB hallmark analysis was performed, revealing elevated IFN I/II signaling signatures in LE skin compared to NL skin, both in expression levels and in the percentage of expressing cells across nearly all cell types and subtypes (Fig. 5I; S6G). Melanocytes showed the most significant upregulation, followed by immune cells and keratinocytes.

Given the considerable overlap in IFN I and IFN II-induced target genes, we investigated ligand expression. We found *IFNG* to be the predominant ligand expressed in NL/LE skin, whereas *IFNA* expression was undetectable, and *IFNB* expression was only detected in DC/MPs (and slightly decreased in LE compared to NL; data not shown). Despite the somewhat reduced *IFNG* expression in immune cells of the LE skin, LE fibroblasts, keratinocytes, and melanocytes all showed increased frequencies of higher *IFNG* expressors compared to NL skin (Fig. 5J). These data suggest the non-immune cells to be the source of elevated IFN signaling in LE skin.

### Small interfering RNA (siRNA)-mediated knockdown of key genes downregulated in LE skin reveals changes in keratinocyte proliferation and stress response, implicating the importance of keratinocytes in VLS pathogenesis

To explore potential causal events in VLS, we turned to *in vitro* monolayer and organotypic models. Specifically, we targeted genes with reduced expression in LE epidermal basal and early suprabasal cells, focusing on three genes identified in our scRNA-seq data – *ATF3*, *APOE,* and *CXCL14* – which showed decreased expression in LE keratinocytes and have largely unknown keratinocyte-intrinsic functions^92,112–114^. We conducted siRNA-mediated knockdown experiments in cultured human N/TERT-2G keratinocytes, an immortalized human keratinocyte cell line^115^.

In monolayer N/TERT-2G cultures, siRNA knockdown of *ATF3* elicited an insignificant proliferation effect, whereas knockdown of *APOE* and *CXCL14* each significantly reduced keratinocyte proliferation (Fig. 6A, B). Remarkably, the expression of stress-associated keratin genes *KRT6/16/17* as well as alarmin genes *S100A8* and *S100A9* was significantly upregulated by *ATF3* or *APOE* deficiency, but not *CXCL14* deficiency (Fig. 6B). To eliminate potential off-target effects, a second siRNA targeting a different region of the cognate gene was used to deplete *ATF3* or *APOE.* In both cases, elevated expression of stress keratin and alarmin genes was observed (Fig. 6C).

**Figure 6.**
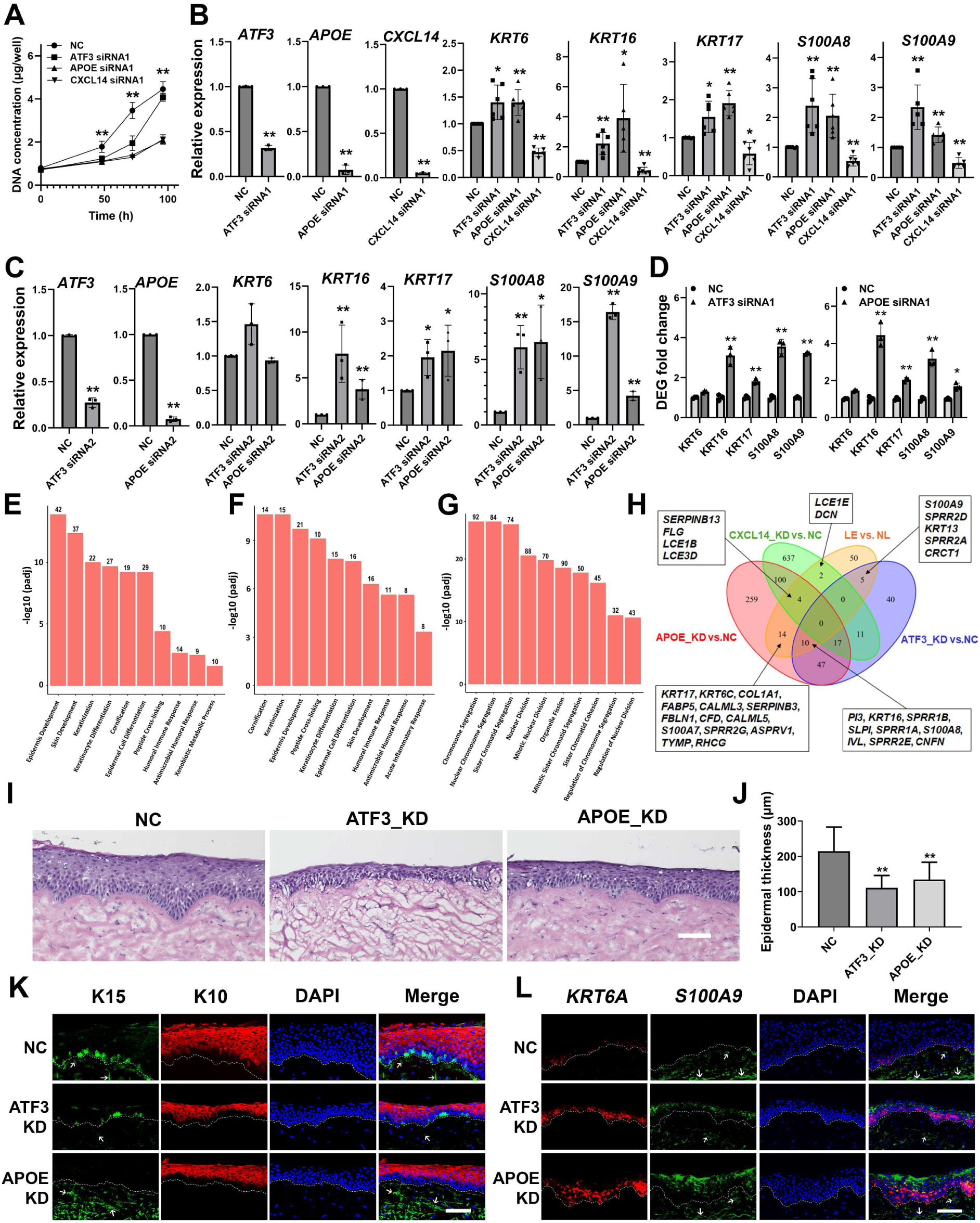
Recapitulation of LS-like keratinocyte stress response upon epidermal-specific depletion of *APOE* and *ATF3*. (A) Effect of siRNAs targeting *ATF3, APOE*, or *CXCL14* on N/TERT-2G keratinocyte proliferation, which was represented by increase in DNA concentration (μg/well) measured at 0, 24, 48, and 96 h after siRNA transfection. NC, scrambled negative control siRNA. (B-C) RT-qPCR results of the indicated genes analyzed at 48 h after siRNA transfection. Note that *ATF3* and *APOE* were each targeted by two different siRNAs (B, #1; C, #2) to minimize off-target effect. (D) Bulk RNA-seq analysis validates the differential expression of the indicated genes. ** *adj p* < 0.01, * *adj p* < 0.05. (E-G) GO Enrichment analysis of the bulk RNA-seq data showing top ten up-regulated biological processes upon knockdown of *ATF3* (E) or *APOE* (F), and top ten down-regulated biological processes upon knockdown of *CXCL14* (G). Number above each bar represents the number of genes with significant changes in the *ATF3, APOE*, or *CXCL14* knockdown (KD) group compared to the NC group. (H) Venn diagram highlighting the overlap of molecular changes across various comparisons: *ATF3* KD vs. NC; *APOE* KD vs. NC; *CXCL14* KD vs. NC; and LE vs. NL in VLS. The numbers of overlapping genes are indicated in the diagram, and gene names are listed inside the rectangles. ** *p* < 0.01, * *p* < 0.05. (I-K) H&E staining (I) of, and epidermal thickness (K) in, 3D-organotypic human skin equivalents derived from N/TERT-2G keratinocytes transfected with NC, *ATF3* (#1), or *APOE* (#1) siRNA. (K-L) Immunofluorescence (K) and RNAScope (L) of the indicated proteins or transcripts in 3D-organotypic human skin equivalents derived from control, *ATF3* KD, or *APOE* KD N/TERT-2G keratinocytes. White arrows indicate autofluorescence in the dermis. Scale bar = 100 μm. ** *p* < 0.01; * *p* < 0.05.

To more comprehensively define the molecular changes induced by the knockdown of these genes, we performed bulk RNA-seq analysis. The results confirm that knockdown of either *ATF3* or *APOE*, but not *CXCL14*, led to elevated expression of stress keratin and alarmin genes (Fig. 6D). GO analysis revealed “epidermal development”, “keratinocyte differentiation”, and “cornification” as top terms that are enriched with *ATF3* or *APOE* knockdown (Fig. 6E, F). Moreover, knockdown of *ATF3* or *APOE* led to enrichment of terms including “humoral immune response” and “antimicrobial humoral response”. These data support an overlapping molecular function of ATF3 and APOE in keratinocyte stress response, epidermal differentiation, and immune response. However, *CXCL14* knockdown predominantly resulted in a downregulation of genes associated with cell cycle progression (Fig. 6G). Importantly, genes upregulated by knockdown of *ATF3*, *APOE*, and *CXCL14* collectively “accounted for” 40% of the upregulated DEGs identified in our scRNAseq data, including stress keratins, alarmins, serine proteinase inhibitor genes, and late differentiation genes (Fig. 6H). These data uncover differential function of the three targeted genes in keratinocyte biology, with *ATF3* and *APOE* being required for maintaining keratinocyte homeostasis and suppressing damage response, and *CXCL14* maintaining the proliferative capacity of keratinocytes.

To investigate the roles of ATF3 and APOE in a more physiological setting that enables epidermal differentiation, we utilized N/TERT-2G-derived 3D skin equivalent organotypic cultures. While epidermal tissue growth and stratification occurred with keratinocytes transfected with negative control siRNA, knockdown of *ATF3* or *APOE* in keratinocytes resulted in the formation of a thinner epidermis (Fig. 6I, J). Importantly, immunofluorescence analysis revealed reduced expression of K15 in the basal layer of knockdown cultures (Fig. 6K), recapitulating the finding in LE skin from VLS patients. Moreover, RNAScope experiments revealed the increased expression of stress-associated keratin (*KRT6A*) and alarmin (*S100A9*) genes in organotypic cultures with *ATF3* or *APOE* knockdown (Fig. 6L). These results identify *ATF3* and *APOE* as likely functional mediators of stress-associated, VLS-like changes in epidermal keratinocytes, and implicate the functional importance of keratinocytes in VLS pathogenesis.

## DICUSSION

Our study highlights VLS-associated molecular alterations across multiple cell types, altered cell-cell interactions, and dysregulated signaling pathways in LE compared to NL skin. Collectively, these findings suggest a multifactorial etiology for VLS, where structural skin cells like fibroblasts, keratinocytes, and melanocytes collaborate with immune cells to induce clinical lesions.

Our data offer an unprecedented overview and provide detailed insights into VLS heterogeneity. At the cellular level, various patterns of deviation were observed in different cell types across LE samples. For instance, some LE samples showed an expansion of CD8^+^ T cells, while others exhibited the presence of B cells. At the molecular level, gene expression and signaling alterations show distinct profiles in individual patients, even when common modules are identified. While the precise mechanism underlying this heterogeneity is unknown, it likely reflects a combination of patient-specific factors, including genetic background and tissue microenvironmental variations, as well as the stage of disease progression (e.g., chronicity and activity). Comprehensive characterization and understanding are crucial for developing precision therapeutic strategies tailored to the specific pathogenic mechanisms of each patient.

Scarring in VLS results in devastating architectural changes of the vulva, compromising sexual and urogenital functions^1^. However, the cellular and molecular basis of dermal scarring in VLS has not been determined. Contrary to expectations for a typical fibrotic disease ^9,116^, our findings indicate a reduced TGF-β signaling potential in the LE skin microenvironment. While the PDGF/PDGFR pathway is linked to proliferative and fibrotic disorders^117,118^, our data indicate reduced PDGF/PDGFR signaling in LE skin, akin to impaired wound healing^119^. LE fibroblasts also lose the basic molecular characteristics of skin fibroblasts, evident through reduced collagen and elastin gene expression. Instead, they upregulate the molecular features associated with an inflammatory function, such as signaling via CXCL12 primarily to T cells, which could play a role in retaining tissue-resident memory T cells in skin. CXCL12 production by a subset of SFRP2^+^ fibroblasts has been found in previous studies to contribute to the amplification of the immune network in psoriasis^120^. Notably, the LE fibroblasts in VLS appear to have reduced potential to signal to typical inflammatory myeloid cells like neutrophils and macrophages through expected cytokines such as CXCL1/2/3 and CCL2. This is seemingly distinct from fibroblasts in other inflammatory skin diseases such as psoriasis, where neutrophil/macrophage involvement is prominent^121,122^. Our detection of reduced expression of *COL6A5* and *COL18A1* in VLS LE fibroblasts is also intriguing, given that a small COL6A5^+^COL18A1^+^ fibroblast subpopulation – characterized as inflammatory because of their expression of inflammatory cytokines such as CCL2 and CCL19 – is enriched in atopic dermatitis lesions^123^. Overall, the molecular landscape in VLS LE fibroblasts cannot be explained by a typical fibrotic phenotype. While early VLS may involve dermal fibrosis, progression to a late, chronic stage results in a degenerative and stressed state. Furthermore, the inflammatory fibroblasts in VLS are likely distinct from those in other skin diseases, influencing specific patterns of inflammation^123^.

The immune alterations detected in LE skin do not provide unequivocal support for a Th1 disease^3,17,72^. On the one hand, we saw evidence of elevated adaptive immunity and increased T cell cytotoxicity in LE skin. The heightened JAK/STAT and IFN signaling activities are also consistent with Th1 signaling being active in LE skin. On the other hand, CD8^+^ T cells, the most predominant source of IFN-γ per our scRNA-seq data, express lower levels of *IFNG* and *TNF* in LE compared to NL skin. These findings, together with the higher expression of *CD69* in NL skin compared to LE skin, suggest that NL skin in VLS patients may already have a prelesional environment with antigen experienced tissue-resident memory T cells. As the disease progresses to clinical lesions, these sentinel memory T cells are likely reactivated and become highly specialized cytotoxic cells that amplify the inflammatory LE skin microenvironment. The presence of B cells in LE skin of some VLS patients further supports an antigen-driven response. Furthermore, our discovery of the CXCL12-CXCR4 signaling axis raises the intriguing possibility that proinflammatory fibroblasts are involved in the retention of tissue-resident memory T cells in VLS skin^124^.

Both IFN-γ and TNF-α are important for activating macrophages^125^. Intriguingly, LE macrophages/DCs, along with CD8^+^ T2 and cycling T cells, express lower levels of *IFNGR1* (IFN-γ receptor) compared to their NL counterparts (Fig. S4H). Moreover, there is a widespread reduction in TNF-α signaling potential within the LE skin microenvironment. Reduced Th1 cytokine production can lead to a weakened immune response against infections and tumors, potentially increasing susceptibility to chronic infections and cancer – a notion consistent with the higher incidence of squamous cell carcinoma in VLS patients^2^. Additionally, while Th1 cytokines are known to promote inflammation, they can also play a regulatory role in immune responses. A reduction in these cytokines may disrupt immune regulation, potentially contributing to the development of autoimmune conditions. Finally, the reduction in transcripts might not reflect the actual cytokine production by T cells activated by antigens, which could be signaling other non-DC cells as their targets to worsen the inflammation. Future studies are needed to understand the implications of these immune changes on the clinical course of VLS.

Our experiments showing keratinocyte cell death and stress response in LE underscore a likely role of keratinocytes in VLS pathogenesis. Tissue damage typically triggers the activation and proliferation of stem and progenitor cells to restore homeostasis, a regenerative response observed across various tissues, including skin, bone, and skeletal muscle^126,127^. However, when cell injury is irreversible, cell death pathways can also be activated^128^. The depletion of stem and progenitor pools, along with premature differentiation of LE keratinocytes, likely reflects this damage-induced, dysregulated tissue regenerative process. The finding that siRNA knockdown of three genes linked to stem/progenitor and early differentiating states can replicate LE-associated gene expression changes suggests that the disruption of progenitor/transitional states creates a self-perpetuating cycle, amplifying tissue damage and leading to atrophy.

Keratinocytes may actively contribute to LE skin inflammation. VLS lesions exhibit elevated potential for annexin A1 signaling, a key factor in Stevens-Johnson Syndrome/Toxic Epidermal Necrolysis that promotes keratinocyte necroptosis^129^. Necroptosis is a recently discovered programmed cell death pathway that, distinct from the inflammation-silent apoptosis, is highly inflammatory through the release of alarmins (e.g., HMGB1 and S1008/9) rather than classical cytokines such as TNF-α^130,131^. Necroptotic keratinocytes in psoriasis-like skin are known to release damage-associated alarmins including S100A8 and S100A9^132^, which can be secreted into the extracellular space and promote the induction of autoreactive CD8^+^ T cells in various autoimmune/inflammatory settings^133^. Elevated expression of S100A8/9 has been detected in lesions of lichen planus and S100A8 expression has been shown to enhance the cytotoxic response of patient CD8^+^ T cells^134^. Moreover, keratinocyte-derived S100A8/9 can activate melanocytes and melanoma cells through paracrine signaling^135^. Therefore, the necroptotic keratinocytes and elevated levels of S100A8/9 in VLS lesions likely influence the vulvar skin microenvironment, affecting CD8^+^ T cells and melanocytes, and thereby promoting disease progression. Given the elevated expression of MHC I and II genes in LE melanocytes, with class II typically expressed by myeloid cells and B cells but rarely by non-immune cells, these melanocytes can become targets for T cells^136^. As a result, T cells can directly kill melanocytes, potentially releasing more antigens and further exacerbating LE pathology.

Our results demonstrate elevated IFN and JAK/STAT signaling in VLS lesions in both clinically early and advanced disease, with JAK/STAT activation seemingly limited to keratinocytes in early disease. Type I IFNs are known to play a pathogenic role in various autoimmune diseases, including systemic lupus erythematosus, systemic sclerosis, dermatomyositis, rheumatoid arthritis, and Sjogren’s syndrome^136,137^. In these settings, type I IFNs enhance antigen presentation, stimulate lymphocyte responses, and trigger the expression of chemokines. Type II IFNs are also implicated in autoimmune and inflammatory disorders^138^. IFN-γ has been shown to enhance cell-mediated cytotoxicity against keratinocytes in other interface-predominant dermatitis, including lichen planus and cutaneous lupus^139,140^. Both types of IFN signaling can induce the expression of MHC I or II genes^141^, and both are known to also exert protective roles in autoimmune diseases^137,138^. The overlap in downstream responsive genes precludes us from determining whether the elevated signaling in LE skin is type I, type II, or both. Plasmacytoid dendritic cells, a major source of type I interferons, have been suggested to be present in VLS skin^142^. However, our scRNA-seq data lacks the resolution to distinguish this DC subset and we did not detect significant levels of IFN-α/β expression in either NL or LE skin. While *IFNG* expression is reduced in CD8^+^ T cells, it is upregulated in fibroblasts, keratinocytes, and melanocytes of LE skin. It remains possible that both type I and II IFN signaling are elevated in LE skin, but with spatiotemporal and cell-type specificities. Regardless, our findings provide a strong foundation for future clinical trials to explore the functional blockade of IFNs and the inhibition of downstream JAK/STAT signal transduction pathways to better understand their roles in VLS pathogenesis, and to evaluate the therapeutic potential of JAK/STAT inhibitors in treating mid-to-late stages of VLS.

In summary, we propose a working model for VLS pathogenesis, where tissue-resident memory T cells are retained and reactivated upon signaling from pro-inflammatory fibroblasts and antigen encounter with melanocytes. Fibroblasts and keratinocytes are not only targets of immune activity but also active contributors to disease by releasing inflammatory cytokines and alarmins that can exacerbate inflammation. While our study significantly advances the understanding of VLS, it is limited in delineating the precise sequence of molecular changes and identifying the cell(s) of origin in the disease. Future research should focus on developing innovative strategies and experimental models to tackle these problems. Such efforts are essential for unraveling the intricate molecular and cellular dynamics that drive VLS initiation and progression, ultimately leading to more effective treatments and interventions.

## MATERIALS AND METHODS

### Study design

This study utilized spatial transcriptomics and scRNA-seq to study LE and NL skin in VLS. Analysis of histologic specimens and punch skin biopsies of patients with a diagnosis of biopsy-proven VLS were performed under IRB-approved protocols at University of California, Irvine (UCI). All samples were de-identified before use in experiments. VLS skin samples were evaluated and obtained by a board-certified dermatologist. Clinically active LS lesions were characterized by the presence of erythema and/or textural change (atrophy, lichenification). Late lesions were characterized clinically by scarring/architectural changes. NL sites were selected as skin adjacent to the site of VLS with the absence of any the morphologic findings (texture change, erythema, hypopigmentation/depigmentation, scarring) of VLS clinically and under dermoscopy.

### Subject enrollment

Five subjects were enrolled for scRNA-seq experiments of both LE and NL skin. Subjects enrolled had not used topical steroids or another topical immunomodulators in the two weeks prior to sample procurement. Patients were not on a biologic or systemic immunomodulator for VLS or another condition. All patients were post-menopausal and were not on topical or systemic hormonal replacement at the time of biopsy. Patients included did not have a diagnosis of a concomitant vulvar dermatoses (e.g. lichen planus) or infectious process.

### Patient samples for immunohistochemistry and immunofluorescence, GeoMx, RNAScope, and Xenium

Human skin FFPE samples, obtained for diagnostic purposes from VLS patients or from benign vulvar excisions in non-VLS patients, were analyzed and subjected to the indicated assays. All studies on FFPE tissue were performed under IRB-approved protocols at UCI.

### Histology, immunostaining, and RNAScope

For paraffin-embedded human vulvar samples, 4-μm sections were stained with hematoxylin and eosin (H&E) using H&E Stain Kit (Abcam) per the manufacturer’s protocol or using the appropriate antibodies following antigen retrieval as previously described^143^. Primary antibodies included: K5, K10, K14, K15, loricrin, CD3, CD45, SMA, vimentin, Ki67, E-cadherin, cleaved caspase-3, and p-MLKL; and fluorescence-conjugated secondary antibodies used included: Alexa Fluor 488-conjugated goat anti-mouse, Alexa Fluor 568-conjugated goat anti-rabbit, Cy5-conjugated donkey anti-mouse, Alexa Fluor 647-conjugated donkey anti-rabbit and FITC-conjugated donkey anti-rabbit (information is provided in Table S7). Immunohistochemical detection was performed using Vector ABC (Vector Laboratories; PK-6100) and DAB (DAKO; K3468) kits according to the manufacturers’ recommendations.

RNAScope was performed as described^144^ using the following probes: Hs-KRT6A, Hs-S100A9, Hs-KRT16, Hs-APOE, Hs-ATF3, and Hs-CXCL14 (information is provided in Table S7). Images were acquired with a Keyence microscope.

Keratinocytes-derived 3D skin organotypic cultures were freshly frozen in Tissue-Tek O.C.T. Compound (Sakura) and sectioned at 10 μm on Superfrost Plus microscope slides (Fisher Scientific). Sections were fixed at room temperature (RT) for 1 hour with 4% paraformaldehyde (Electron Microscopy Sciences) diluted in 1x PBS (Gibco), followed by H&E staining or immunofluorescence using the indicated primary antibodies in conjunction with Alexa Fluor 488-conjugated goat anti-mouse and Alexa Fluor 568-conjugated goat anti-rabbit as secondary antibodies (Table S7).

### GeoMx spatial transcriptomics

FFPE specimens were sectioned and mounted onto histology-grade glass slides. Slides were prepared following GeoMx-NGS Slide Preparation User Manual (MAN-10115-05) following the RNA FFPE Manual Slide Preparation Protocol section in the user manual. Briefly, slides were deparaffinized using Xylene, subjected to antigen retrieval, digested with proteinase K, hybridized overnight with Human Whole Transcriptome Atlas probe set (Cat# 121401102), and stained with morphology markers (pan-cytokeratin, SMA, and Syto83). The hybridized and stained slides were loaded onto the GeoMx instrument following GeoMx-NGS DSP Instrument User Manual (SEV-00087-05). Collected ROIs aspirate was processed following GeoMx-NGS Readout Library Prep User Manual (MAN-10117-05). NGS library was sequenced on the Illumina NovaSeq 6000 platform, targeting 200 reads per um^2^.

FASTQ files were converted to DCC files using GeoMx NGS Pipeline software. Briefly, reads were aligned to RTS-ID barcode list, PCR duplications were removed, then converted to DCC files. GeoMx data was transformed into a Seurat object for analysis. Gene counts were Q3 normalized considering all target values, and 3000 informative features were selected. Integration was then performed using the FindIntegrationAnchors and IntegrateData functions. The top 30 principal components (PCs) with a resolution of 0.5 were used to identify cell clusters. FindAllMarker function with default parameters was used to find marker genes for each cluster.

### scRNA-seq

Generation of single-cell suspensions was performed as previously described^145^. Briefly, samples were incubated in 5 U/mL dispase (StemCell, Cat# 07913) at 37℃ for 45 min. The epidermis was peeled from the dermis, cut into fine pieces, and incubated in 0.25% trypsin-EDTA (Gibco, Life technologies, Cat# 25200) with 50 μg/mL DNase I (Thermo Fisher Scientific, Cat# DN25-100M) for 45 min at 37°C, quenched with fetal bovine serum, and passed through a 70-μM cell strainer. Dermis was minced and digested in 0.1% collagenase II (Life Technologies, Cat# 17101-015) and 0.1% collagenase V (Sigma, Cat#C9263) in RPMI 1640 with 50 μg/mL DNase I for 90 min at 37°C and strained through a 70-μM cell strainer. Dead cell removal was performed using a Dead Cell Removal Kit (Miltenyi Biotec). Epidermal and dermal cells were then recombined and washed in PBS containing 0.04% BSA and resuspended at a concentration of approximately 1,000 cells/µL.

Library generation was performed following the 10x Genomics Chromium Single Cell 3ʹ v3.1 Reagents Kit (following the CG000315 Rev E. user guide), where we targeted 10,000 cells per sample for capture. Additional reagents included: nuclease-free water (Thermo Fisher Scientific; AM9937), low TE buffer (Thermo Fisher Scientific; 12090-015), ethanol (Millipore Sigma; E7023-500ML), SPRIselect Reagent Kit (Beckman Coulter; B23318), 10% Tween 20 (Bio-Rad; 1662404), glycerin (Ricca Chemical Company; 3290-32), Qiagen Buffer EB (Qiagen; 19086). Each library was sequenced on the Illumina NovaSeq 6000 platform to target an average of approximately 50,000 reads per cell.

FASTQ files were aligned utilizing 10x Genomics Cloud Cell Ranger Count v7.1.0. Each library was aligned to an indexed Human GRCh38 genome. Preliminary analysis and visualization of the single-cell datasets were performed using the Seurat R package version 4.3.1. Low-quality and dead cells (nGenes <500 or >6,000 or percent mitochondria > 10%) were excluded from the analysis. Following SCTransform normalization of each patient dataset, 3000 informative features were selected and all eight datasets were integrated using the FindIntegrationAnchors function. This was followed by PC analysis (PCA), generation of Elbow Plot, and UMAP analysis using the first 20 PCs, RunUMAP and FindNeighbors. Doublets were removed in two steps. All cells were first clustered with a low resolution (R) of 0.2 to remove large doublet clusters. Clustering at a high R of 6 was then performed, resulting in numerous small clusters that enabled the maximal removal of doublets. Subsequent analysis was using an R of 0.3.

For sub-clustering analysis, we computationally isolated all cell clusters that expressed *PDGFRA* for fibroblasts, *PTPRC* for immune cells, and keratin markers *KRT5* and *KRT1* for keratinocytes from the integrated master dataset (Fig. 3B). Downstream subclustering analysis was performed using R of 0.1 for fibroblasts, 0.25 for immune cells, and 0.16 for keratinocytes, using PC of 20 for all.

### Cell-cell communication and RNA velocity analyses

For cell-cell communication predictions, CellChat analysis was performed for each condition of the log-normalized integrated data for all patients, or pair-wise data for each individual patient. The default parameter settings were adopted except that the ligand-receptor interaction database was set to be “Secreted Signaling”. A total of 20 cell clusters were used for analysis, including six keratinocyte clusters (BSC, BC, SC1, SC2, GC, and cycBC), seven immune cell clusters (CD4T1, CD4T2, CD8T1, CD8T2, cycTC, DC/MP, and MC), three fibroblast clusters (FB1, FB2, and FB3), smooth muscle cells (SM), pericytes (PC), endothelial cells (EC), and melanocytes (ME).

Three methods, TFvelo, scVelo and nlvelo, were used for RNA velocity analysis on keratinocytes under each condition, using default parameter settings. Both scVelo and nlvelo require the spliced and unspliced RNA counts, with scVelo modeling the splicing dynamics using linear equations and nlvelo using non-linear equations^146^. TFvelo estimates transcriptional-regulation-driven RNA velocity by modeling the dynamics of target genes and their transcription factors, solely based on the single-cell RNA counts without relying on splicing information^147^. For scVelo and nlvelo, loom files that contain the spliced and unspliced RNA counts were generated using velocyto package (version 0.17). For nlvelo and TFvelo, 1,000 cells from each condition were randomly sampled for analysis to save time.

### Gene scoring, GO, and PROGENy analyses

Gene scoring analysis was performed in AddModuleScore function using appropriate gene lists as previously described^146^.

GO enrichment for biological processes was done by selecting the markers (-0.5 ≥ log2FC ≥ 0.5; padj < 0.05) identified in ‘FindMarkers’ and executing the ‘enrichGO’ function of ‘clusterProfiler’ (Version 4.8.3) on that dataset using the ‘org.Hs.eg.db’ database. The GO terms were presented by plot using ggplot2 (Version 3.4.4). GO terms with a *p*-value < 0.05 were considered as significantly enriched. Analysis of fibroblast and immune cell datasets was done after removing keratinocyte-specific genes from the DEG lists (Table S4 and S5).

Pathway activity scores were calculated using the ‘progeny’ function (organism = “Human”, top = 100) from ‘PROGENy’ (v1.22.0).

### Xenium spatial transcriptomics

FFPE specimens were sectioned and mounted onto Xenium slides (PN-1000465) following Xenium In Situ for FFPE – Tissue Preparation Guide (CG000578 Rev C). Slides were deparaffinized and decrosslinked following Xenium In Situ for FFPE – Deparaffinization & Decrosslinking (CG000580 Rev C). Immediately, slides were processed following Xenium In Situ Gene Expression with Cell Segmentation Staining (CG000749 Rev A). Briefly, probe hybridization was performed overnight with 10x predesigned Human Skin panel (Cat# 1000643) supplemented with 100 add-on custom probes (information will be provided upon request). Tissues were stained overnight with the Xenium Multi-tissue stain kit, followed by immediate loading onto the Xenium analyzer per Xenium Analyzer manual (CG000584 Rev E).

For data analysis, only cells with at least 20 transcripts and genes detected in at least one cell were retained for further analysis. Data integration was performed using SCTransform workflow with default parameters, followed by dimensionality reduction and clustering using Seurat commands RunPCA, RunUMAP, FindNeighbors, and FindClusters. For RunUMAP and FindNeighbors, the first 20 PCs were used. The resolution parameter for FindClusters was set to 1.5. The clusters were combined based on marker expression, which identified the types of clusters. A cell group comma-separated value (CSV) file was created and uploaded to Xenium Explorer.

### Keratinocyte culture and siRNA transfection

N/TERT-2G human keratinocytes were kindly provided by Jos P. H. Smits (Radboud University Medical Center, Nijmegen, The Netherlands), with permission from Dr. James Rheinwald (Harvard Medical School)^148^.

Stealth duplex siRNAs targeting *ATF3* (Gene bank ID: NM_001674.4), *APOE* (Gene bank ID: NM_001302691.2), and *CXCL14* (Gene bank ID: NM_004887.5) were designed using Invitrogen BLOCK-iT RNAi Designer and were synthesized commercially (Thermo Fisher Scientific). Sequences of all the siRNAs are listed in Table S7. A stealth negative control siRNA with 48% guanine and cytosine content (Thermo Fisher Scientific) was used as a scrambled control.

For siRNA knockdown, N/TERT-2G keratinocytes were plated on 6-well plates (Falcon, 40,000 cells per well) or 24-well plates (Falcon, 15,000 cells per well) in Keratinocyte SFM (1X) medium (Gibco), and cultured at 37°C with 95% air/5% CO_2_. At 40-50% confluency, cells were transfected with 100 pmol/well (6-well plate) or 20 pmol/well (24-well plate) of ATF3 siRNA, APOE siRNA, CXCL14 siRNA, or negative control siRNA, using 5 μl/well (6-well plate) or 1 μl/well (24-well plate) of Lipofectamine 2000 transfection reagent (Thermo Fisher Scientific), following the manufacturer’s protocol. Six hours later, the medium was replaced with Keratinocyte SFM (1X) medium supplemented with Penicillin-Streptomycin (Gibco, 100 units/ml). Cells were harvested at 0, 24, 48, and 72 hours post-transfection for proliferation analysis, or at 48 hours for RT-qPCR or use in 3D skin equivalent organotypic culture.

Proliferation analysis was performed based the method of McFarland laboratory^149^. Plates (24-well) of keratinocytes were frozen overnight at -80°C, and thawed at RT for 15 min. Following trypsinization with 200 µl/well of 0.05 % trypsin-EDTA (Gibco) in TNE buffer (10 mM Tris, 2 M NaCl, and 1 mM EDTA) at RT for 7 min, the plates were frozen-thawed one more time. 1.8 mL of 0.2% Hoechst 33258 (Thermo Fisher Scientific) in TNE buffer was added to each well, and the plates were gently agitated for 2 hours at RT. Hoechst 33258 intensity was measured using a Varioskan LUX multimode microplate reader (Thermo Fisher Scientific) with an excitation wavelength of 355 nm and an emission wavelength of 460 nm. A standard curve was constructed using double-stranded calf thymus DNA (Sigma-Aldrich) to determine the sample DNA concentration. The proliferation analysis was repeated in three independent cultures with four replicate wells per treatment per culture.

### RT-qPCR and bulk RNA-seq

Total RNAs were isolated using the Quick-RNA™ Miniprep Kit (Zymo Research) following the manufacturer’s protocol. Equal amounts of RNA from each treatment group were reverse transcribed into cDNA using the High-Capacity cDNA Reverse Transcription Kit (Applied Biosystems). RT-qPCR was conducted on a CFX96 Real-Time PCR Detection System (Bio-Rad) using iTaq Universal SYBR Green Supermix (Bio-Rad). Comparative analysis was performed using the ΔΔCt method to compare the expression of target genes to the housekeeping gene GAPDH. The sequences of all RT-qPCR primers are provided in Table S7.

Total RNAs were sent to Novogene Corporation Inc. (Sacramento, CA, USA) for bulk RNA-seq analysis. Briefly, mRNA was purified from total RNA using poly-T oligo-attached magnetic beads. After fragmentation, the cDNA was synthesized using random hexamer primers. The library was checked with Qubit and real-time PCR for quantification and bioanalyzer for size distribution detection. Quantified libraries were pooled and sequenced on Illumina platforms. Raw data were processed to remove reads containing adapter, and remove reads containing ploy-N (>10% uncertain nucleotides) and low-quality reads (>50% of the bases with quality score <5). All the downstream analyses were based on these high-quality clean data. Reference genome (hg38) and gene model annotation files were downloaded from UCSC genome database. Index of the reference genome was built using Hisat2^150^ v2.0.5 and paired-end reads were aligned to the reference genome using Hisat2 v2.0.5. featureCounts^151^. v1.5.0-p3 was used to count the reads numbers mapped to each gene. FPKM (Fragments Per Kilobase of transcript sequence per Millions base pairs) values for each gene were calculated based on the length of the gene and the read count mapped to it. Differential expression^152^analysis of the three pairs of comparisons (NC vs. *ATF3* KD, NC vs. *APOE* KD, and NC vs. *CXCL14* KD), each with three biological replicates, was performed using the DESeq2 package (1.20.0) in R (4.0.0). The *p*-values were adjusted using the Benjamini and Hochberg’s approach for controlling the false discovery rate. Genes with an adjusted *P*-value ≤ 0.05 were considered differentially expressed. GO enrichment analysis of DEGs was implemented by the clusterProfiler (4.6.0) R package. GO terms with corrected *p*-values < 0.05 were considered significantly enriched.

### Human skin equivalent organotypic culture

Split-thickness human skin was purchased from the New York Firefighters Skin Bank. Devitalized de-epidermal human dermis (DED) was prepared following the protocol by Li and Sen, 2015^153^. The DED was cut into 1.5 cm x 1.5 cm pieces, washed twice in complete DMEM (DMEM supplemented with 10% fetal bovine serum and 100 units/ml penicillin-streptomycin), and incubated in complete DMEM at 37°C overnight before use. For the assembly of organotypic tissue, the cut DED was stretched in a 12-well plate, ensuring the basement membrane was oriented downward and attached to the well bottom. Human primary dermal fibroblasts were isolated from discarded neonatal foreskin, as described previously by Lutz-Bonengel et al. in 2021^154^ and maintained in complete DMEM. The primary fibroblasts (7.5 x 10⁵) were seeded on top of the DED in 1.5 ml of complete DMEM and the resulting cultures incubated at 37 °C for 2 days. Subsequently, the fibroblast-containing DED was inverted and transferred to a new well of 12-well plate with the basement membrane facing up. The culture medium was switched to K-SFM, and N/TERT-2G cells (7.5 x 10⁵) were seeded onto the basement membrane side in a total volume of 1.5 ml. After 2 days, the DED culture was moved into the upper chamber of a 40-μm cell strainer placed within a 6-well tissue culture plate filled with 4 ml of 3dGRO differentiation medium (Millipore Sigma), allowing the basement membrane side to be exposed to air. Differentiation medium was changed every other day, and organotypic tissue harvested after 14 days, freshly frozen in O.C.T. embedding compound on dry ice, and stored at -80°C until use.

## Data availability

The sequencing data reported in this work have been deposited in the GEO database under accession GSE274837 (scRNA-seq) and GSE274068 (RNA-seq). Information required to reanalyze the data reported in this paper is available from the lead contact upon request.

## ACKNOWLEDGEMENTS

We thank the Genomics Research and Technology Hub at the University of California, Irvine (UCI) for expert service. This work was supported by UCI institutional seed grants from Institute for Precision Health, Skin Biology Resource-Based Center (P30-AR075047), Office of Research, and School of Medicine Department of Dermatology. Research personnel were partially supported by NIH Grants R35 GM145307 (X.D.), U01-AR073159 (Q.N., X.D.), NSF Grants DMS1763272 (Q.N., X.D.), Simons Foundation Grant 594598 (Q.N., X.D.), and CBET Grant 2134916 (S.X.A.). J. X. has been supported by the California Institute of Regenerative Medicine Training Grant (EDUC4-12822). S.S. was supported by the NIH T32 Interdisciplinary Training Program in Skin Biology (AR080622).

## AUTHOR CONTRIBUTIONS

X.D. and C.K. conceived the study and directed the project. P.S., C.K., J.X., S.S., Q.N., and Y.J. conducted the experiments. P.S., C.K., W.Z., and A.N. performed the computational analysis. M.O. and R.T. provided guidance on transcriptomics and immune cell analyses, respectively. J.S. provided reagents and guidance for stressed keratinocyte and melanocyte experiments. S.A. and B.S. supervised the organotypic culture experiments. A.E. provided dermatopathology support. Q.N. provided guidance on CellChat analysis. C.K. and X.D. wrote the manuscript with input from all authors.

