## Supplemental Figures and Table 1 for "Single-cell and spatial transcriptomics of vulvar lichen sclerosus reveal multi-compartmental alterations in gene expression and signaling cross-talk"

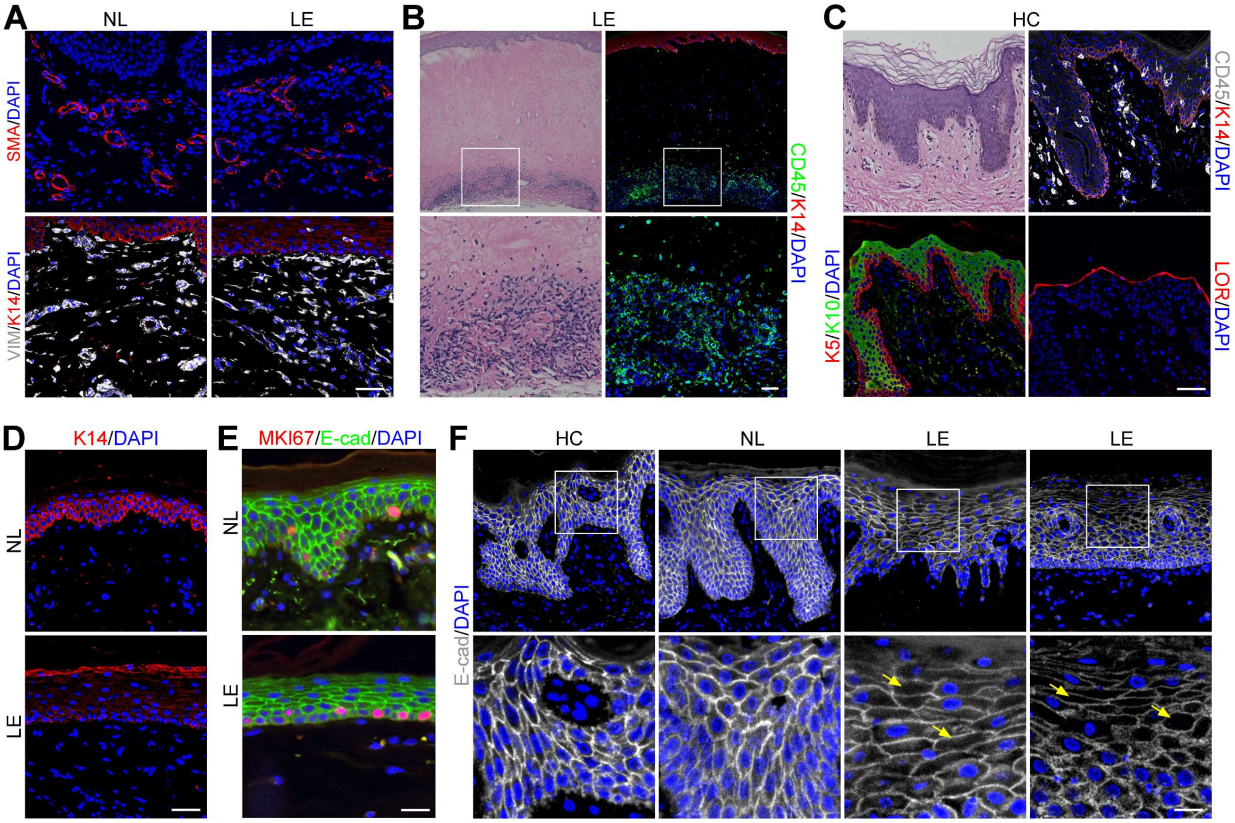


**Figure S1. Additional histological and immunofluorescence analyses of control (HC and NL) and LE skin samples.** Related to Figure 1. (A) Immunofluorescence for SMA (top), vimentin, and K14 (bottom). Scale bar = 50 µm. (B) H&E (left) and CD45/K14 immunostaining (right) of a sclerotic LE sample with cellular aggregates deep in the reticular dermis. Bottom images represent a higher magnification of the boxed areas in top images. Scale bar = 100 µm and 27 µm in top and bottom panels, respectively. (C) H&E and immunostaining analysis of HC control. Scale bar = 50 µm. (D) LE sample from a VLS patient shows altered K14 expression compared to matched NL sample (bottom). Scale bar = 50 µm. (E-F) Ki67 (E) and E-cadherin (F) immunostaining HC, NL and LE samples. Bottom images in F represent a higher magnification of the boxed areas in top images. Scale bar = 25 µm in E; 50 µm and 15 µm in top and bottom panels of F, respectively.

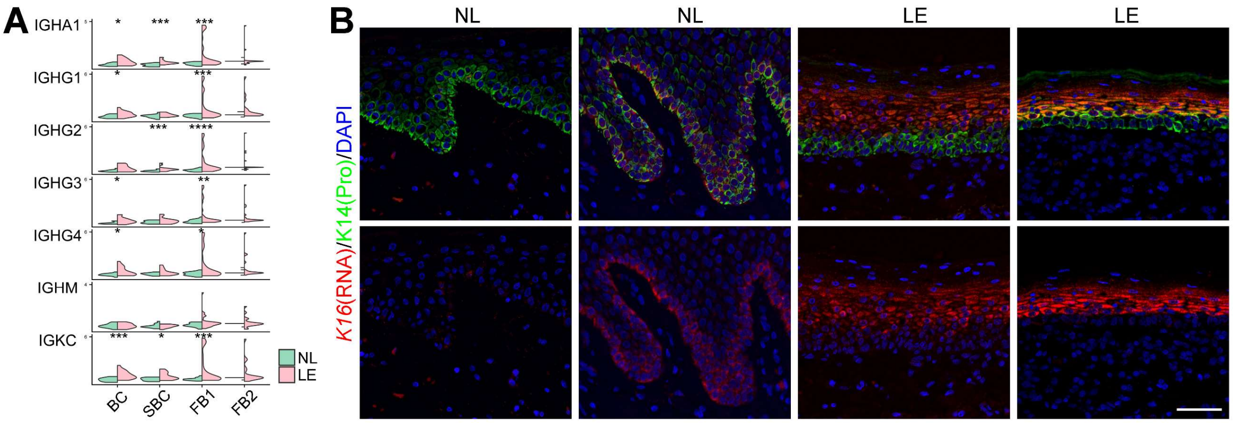


**Figure S2. GeoMx and RNAScope data revealing *IgG* and *KRT16* expression in LE samples.** Related to Figure 2. (A) *IgG* gene expression by cluster in NL compared to LE skin by GeoMx analysis. BC, basal cells; SBC, suprabasal cells; FB, fibroblasts. * $p < 0.05$, ** $p < 0.01$, *** $p < 0.005$, **** $p < 0.001$. (B) RNAScope data showing spatial distribution of *KRT16* transcripts along with K14 protein in LE compared to NL skin. Scale bar = 50 μm.

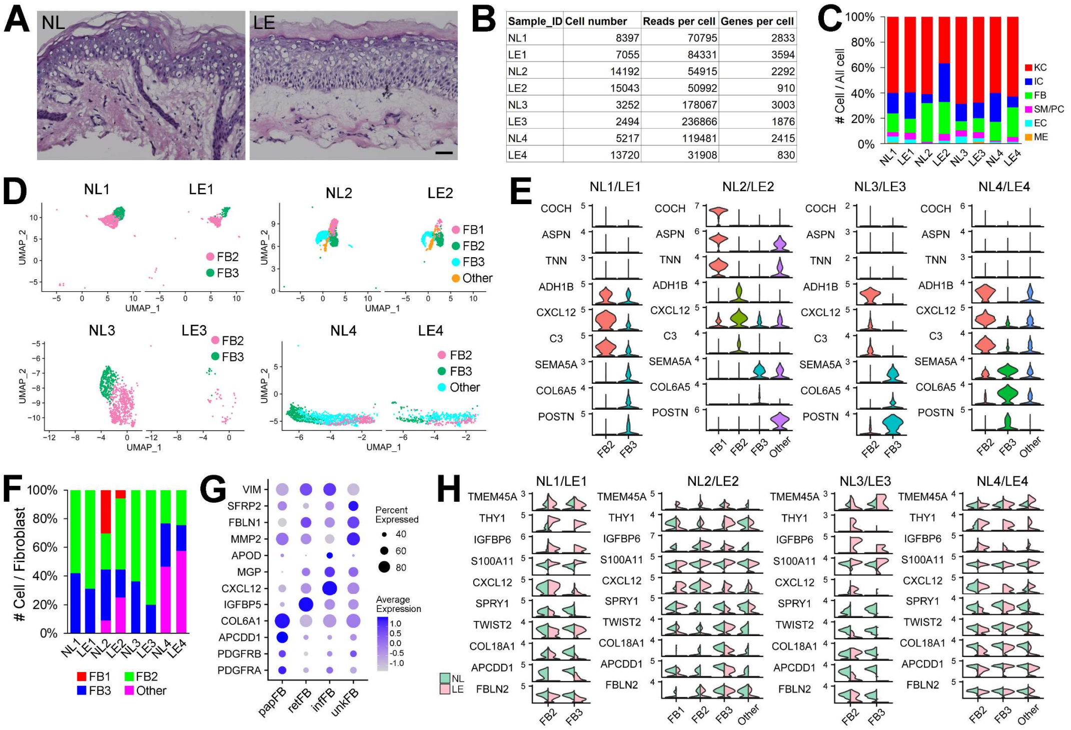


| Sample_ID | Cell number | Reads per cell | Genes per cell |
|---|---|---|---|
| NL1 | 8397 | 70795 | 2833 |
| LE1 | 7055 | 84331 | 3594 |
| NL2 | 14192 | 54915 | 2292 |
| LE2 | 15043 | 50992 | 910 |
| NL3 | 3252 | 178067 | 3003 |
| LE3 | 2494 | 236866 | 1876 |
| NL4 | 5217 | 119481 | 2415 |
| LE4 | 13720 | 31908 | 830 |

**Figure S3. Overall sequencing information for scRNA-seq, with pairwise analyses of fibroblast subclusters from NL and LE samples per patient**. Related to Figure 3. (A) Histology of NL and LE skin from patient 5. Scale bar = 25 µm. (B) Table with number of captured cells per skin biopsy sample, average reads per cell, and genes per cell. (C) Bar plot showing relative proportions of each of the major cell types per patient and sample identities in the integrated dataset. KC, keratinocytes; IC, immune cells; FB, fibroblasts; SM/PC, smooth muscle cell/pericytes, EC, endothelial cells; ME, melanocytes. (D) UMAPs of fibroblast subclusters using pairwise aggregation of patient-matched NL and LE samples without batch correction. Patient 1 = NL1, LE1; Patient 2 = NL2, LE2; Patient 3 = NL3, LE3; Patient 4 = NL4, LE4. (E) Violin plots of marker genes for fibroblast subclusters in each patient. (F) Bar plot showing relative proportions of each fibroblast subcluster using pairwise aggregation. (G) Dot plot from Xenium analysis showing expression of the indicated marker genes to facilitate fibroblast subset identification. papFB, papillary fibroblasts; retFB, reticular fibroblasts; infFB, inflammatory fibroblasts; unkFB, fibroblasts of unknown identity. (H) Split violin plots of select top DEGs in fibroblast subclusters identified using pairwise aggregation.

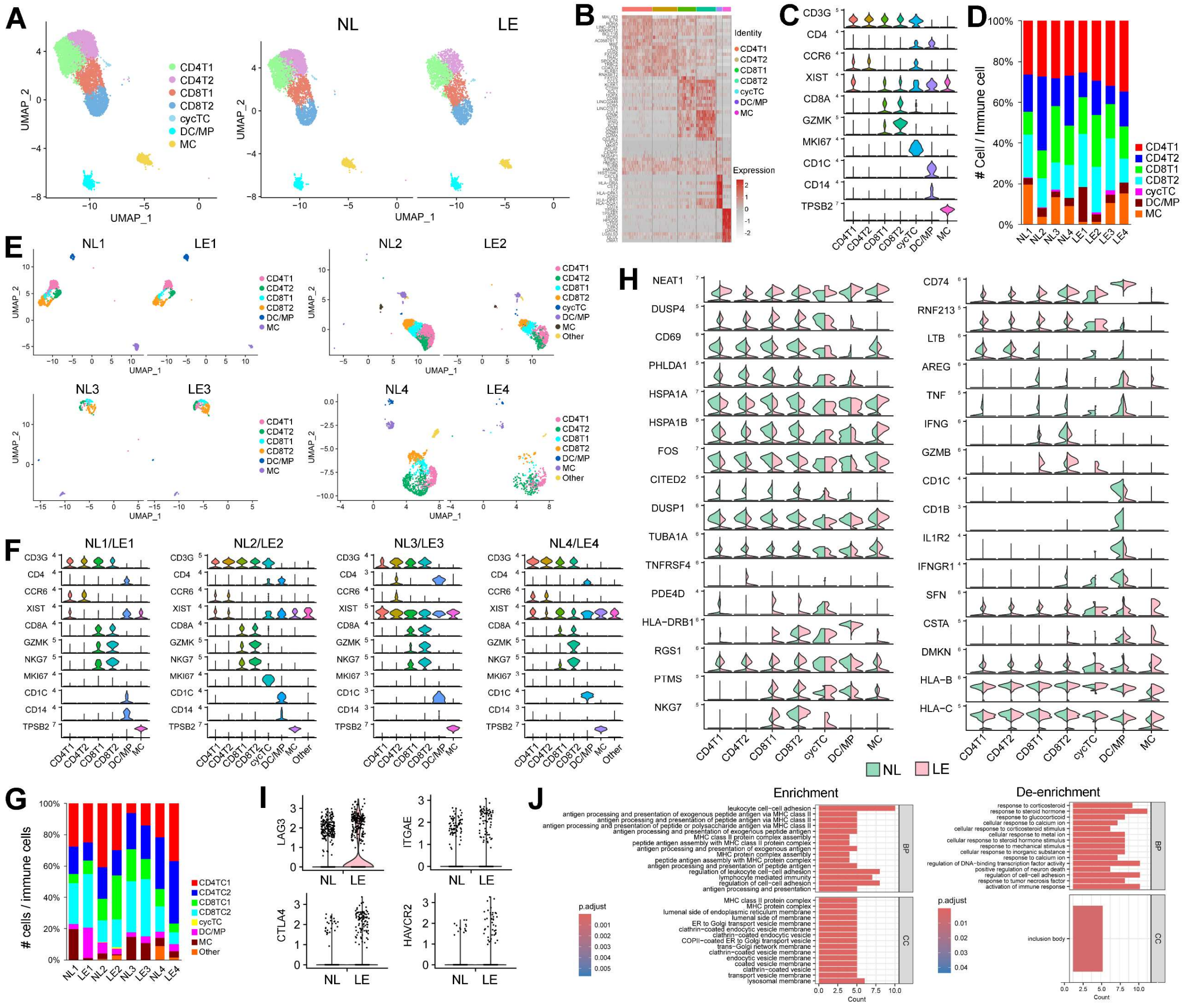

**Figure S4. Molecular alterations in LE immune cells.** Related to Figure 3. (A) UMAPs of immune cell subclusters from the integrated dataset. CD4T1, $CD4^+$ T cell 1; CD4T2, $CD4^+$ T cell 2; CD8T1, $CD8^+$ T cell 1; CD8T2, $CD8^+$ T cell 1, cycTC, cycling T cell; DC/MP, dendritic cell/macrophage; MC, mast cell. (B) Heatmap with top ten marker genes for each subtype of immune cells. (C) Violin plots of select markers for each subtype of immune cells. (D) Bar plot showing relative proportions of each of the immune cell populations per patient and sample identities. (E) UMAPs of immune cell subclusters using pairwise aggregation of patient-matched NL and LE samples without batch correction. (F) Violin plots of select markers for each subtype of immune cells using pairwise aggregation. (G) Bar plot showing relative proportions of each immune cell subcluster using pairwise aggregation. (H) Split violin plots of select top DEGs in immune cell subclusters identified using pairwise aggregation. (I) Violin plots of *LAG3*, *ITGAE*, *CTLA4*, and *HAVCR2* expression in CD8T2 population in NL compared to LE. (J) Diagram showing GO biological process terms enriched (left) and de-enriched (right) in LE immune cells compared to NL immune cells.

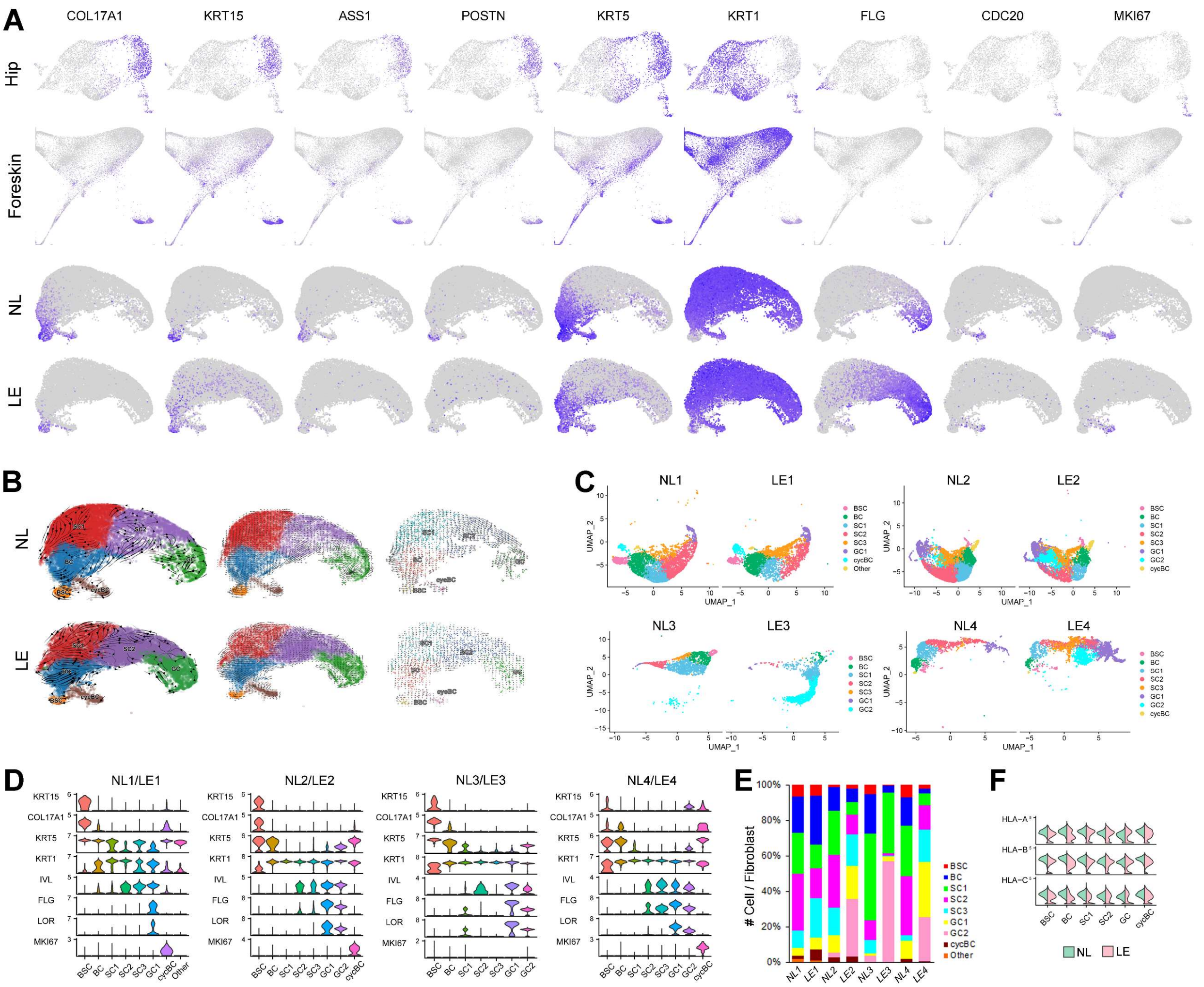


**Figure S5. Epidermal organization in vulvar skin compared to other skin sites, velocity analyses, and pairwise analysis of keratinocytes.** Related to Figure 4. (A) Feature plots of the indicated genes, including epidermal differentiation markers, in hip and foreskin in published data (top panels) and in integrated scRNA-seq data from vulvar keratinocytes in NL and LE skin (bottom panels). (B) Projection of RNA velocity fields onto the UMAP space of keratinocytes in NL and LE from integrated data using two different methods: scVelo (Streamlines, left; Fine-grained, middle) and nlvelo (Fine-grained, right). (C) UMAPs of keratinocyte subclusters using pairwise aggregation of patient-matched NL and LE samples without batch correction. (D) Violin plots of select markers for each subtype of keratinocytes using pairwise aggregation. (E) Bar plot showing relative proportions of each keratinocyte subcluster using pairwise aggregation. (F) Split violin plots of *HLA-A/B/C* expression in keratinocyte subclusters from the integrated data.

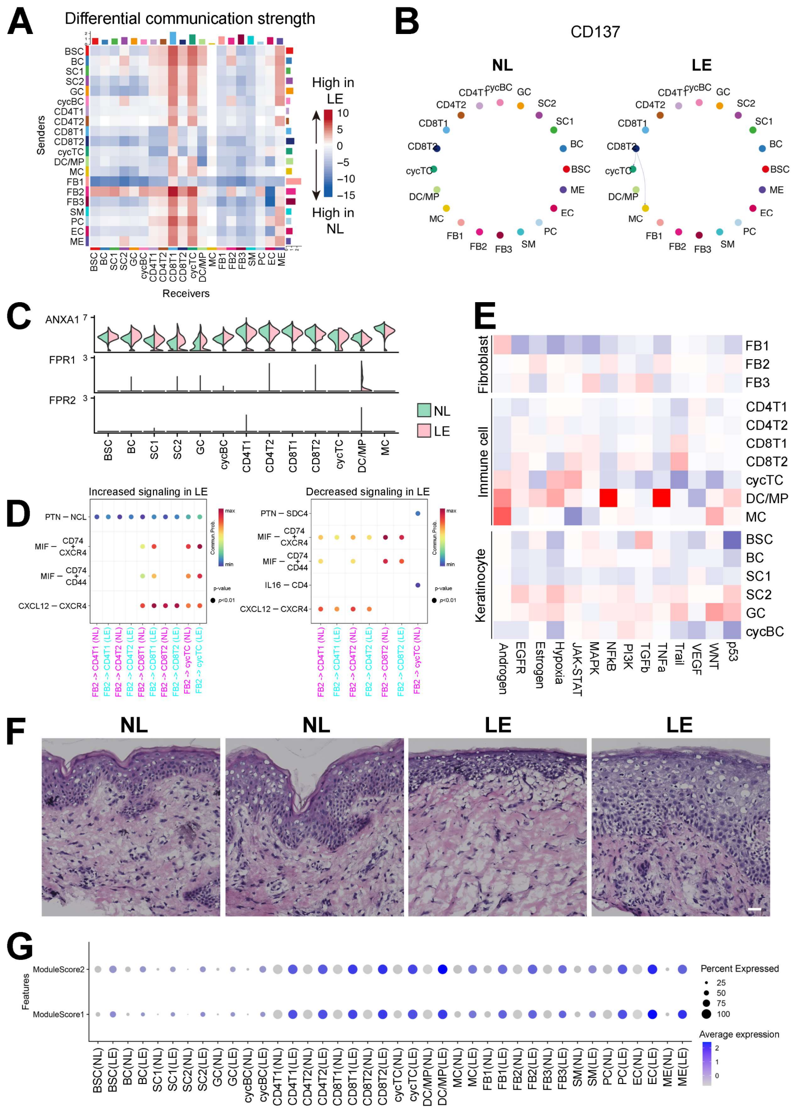


**Figure S6. Inferred alterations in signaling network structure, strength, and downstream activity in LE skin.** Related to Figure 5. (A) CellChat analysis following removal of most of the fibroblasts contaminated with keratinocyte transcripts, showing heatmap (by weight) summarizing differential communication strength between any two cell groups in LE compared to NL. See Fig. 5B for comparison. (B) Chord diagrams of the inferred CD137 signaling network in NL vs. LE skin. (C) Violin plots of expression of *ANXA1* and its receptors, *FRP1* and *FRP2*, in all keratinocyte and immune cell subclusters. (D) Dot plots showing the inferred alterations in FB2-to-immune cell signaling for the indicated pathways in LE. (E) PROGENy analysis using scRNA-seq data of signaling pathway activation by cell type/subtype in fibroblasts, immune cells, and keratinocytes. (F) Histological images of two different regions of NL and LE skin from patient 4. Scale bar = 25 µm. (G) Dot plot of gene signatures using MSigDB for IFNA (ModuleScore1) and IFNG (ModuleScore2) for all cell type/subtypes in NL compared to LE. The size of the dot represents the percentage of expressing cells within each cell type, while the color indicates the average expression level (blue is high).

### SUPPLEMENTAL TABLES

**Table S1**. Histologic features of VLS with subtype designation based on degree of inflammation and sclerosis. Related to Figure 1.

| Histologic Features | n | % |
|---|---|---|
| Sclerosis | 17 | 81 |
| Brisk lymphocyte infiltration (dermis) | 11 | 52 |
| Hyperkeratosis | 9 | 43 |
| Spongiosis | 7 | 33 |
| Lichenoid interface | 7 | 33 |
| Effacement of rete | 6 | 29 |
| Subepidermal split | 5 | 24 |
| Papillary edema | 4 | 19 |
| Epidermal atrophy | 3 | 14 |
| Basal keratinocyte necrosis/apoptosis | 3 | 14 |
| Sparse lymphocyte infiltration (dermis) | 3 | 14 |
| Melanophages | 2 | 10 |
| Moderate lymphocyte infiltration (dermis) | 2 | 10 |
| Hypergranulosis | 2 | 10 |
| Lymphocytes in epidermis | 1 | 5 |
| Dermal fibrosis | 1 | 5 |

| Histological Subtype | n | % |
|---|---|---|
| Inflammatory, non-sclerotic | 4 | 19 |
| Inflammatory (mod-brisk), sclerotic | 10 | 48 |
| Inflammatory (sparse), sclerotic | 3 | 14 |
| Non-inflammatory, sclerotic | 3 | 14 |
| Non-inflammatory, dermal fibrosis | 1 | 5 |

*See Attached Excel Files for "Table S2-S7", and Table titles below.*

**Table S2**. Marker genes of the GeoMx/Seurat-identified basal, suprabasal, FB1, and FB2 clusters in the integrated NL and LE dataset. Top differentially expressed genes in basal, suprabasal, FB1, and FB2 cells between NL and LE samples. Related to Figure 2.

**Table S3.** Clinical characteristics of VLS patients studied in scRNA-seq analysis. Marker genes for all cell types in the integrated NL and LE samples for scRNA-seq analysis. Related to Figure 3. Patients had not used topical or systemic steroids for at least two weeks prior to biopsy.

**Table S4.** Marker genes for all fibroblast clusters in the integrated NL and LE samples for scRNA-seq analysis. Top differentially expressed genes for FB1, FB2, and FB3 subpopulations between NL and LE samples in scRNA-seq analysis. List of keratinocyte transcripts removed from fibroblast analysis. Related to Figure 3.

**Table S5.** Marker genes for the immune cell clusters in the integrated NL and LE samples for scRNA-seq analysis. Top differentially expressed genes for CD4T1, CD4T2, CD8T1, CD8T2, cycTC, DC/MP, and MC cells between NL and LE samples. List of keratinocyte transcripts removed from immune cell analysis. Related to Figure 3.

**Table S6.** Marker genes for the keratinocyte clusters in the integrated NL and LE samples for scRNA-seq analysis. Top differentially expressed genes for BSC, BC, SC1, SC2, GC, and cycBC between NL and LE samples. Related to Figure 4.

**Table S7.** Information for siRNAs, RT-qPCR primers, RNAScope probes, primary and secondary antibodies.
